# On-microscope staging of live cells reveals changes in the dynamics of transcriptional bursting during differentiation

**DOI:** 10.1101/2021.11.26.470114

**Authors:** D.M. Jeziorska, E.A.J. Tunnacliffe, J.M. Brown, H. Ayyub, J. Sloane-Stanley, J.A. Sharpe, B.C. Lagerholm, C. Babbs, A.J.H. Smith, V.J. Buckle, D.R. Higgs

## Abstract

Determining the mechanisms by which genes are switched on and off during development and differentiation is a key aim of current biomedical research. Gene transcription has been widely observed to occur in a discontinuous fashion, with short bursts of activity interspersed with longer periods of inactivity. It is currently not known if or how this dynamic behaviour changes as mammalian cells differentiate. To investigate this, using a newly developed on-microscope analysis, we monitored mouse α-globin transcription in live cells throughout sequential stages of erythropoiesis. We find that changes in the overall levels of *α*-globin transcription are most closely associated with changes in the fraction of time a gene spends in the active transcriptional state. We identify differences in the patterns of transcriptional bursting throughout differentiation, with maximal transcriptional activity occurring in the mid-phase of differentiation. Early in differentiation, we observe increased fluctuation in the patterns of transcriptional activity whereas at the peak of gene expression, in early and intermediate erythroblasts, transcription appears to be relatively stable and efficient. Later during differentiation as *α*-globin expression declines, we again observed more variability in transcription within individual cells. We propose that the observed changes in transcriptional behaviour may reflect changes in the stability of enhancer-promoter interactions and the formation of active transcriptional compartments as gene expression is turned on and subsequently declines at sequential stages of differentiation.

## Introduction

Precise spatio-temporal transcriptional control of gene expression is required to accurately produce developmental changes within a tissue or organism. Mis-regulation of this process is frequently associated with inherited and acquired genetic disease and highlights the importance of understanding in detail how transcription is regulated. While bulk and single-cell sequencing have advanced our understanding of transcription during both normal and abnormal development, these methods only provide a static view of transcription and mRNA abundance. Observing real-time fluctuations in nascent transcription, in individual cells throughout lineage specification and differentiation, is required to fully understand the mechanistic details of gene expression (Brouwer and Lenstra, 2019). Establishing when and how transient and dynamic gene activation occurs at different times during lineage-specification, differentiation, and development is therefore of considerable current interest.

Methods allowing the visualisation of nascent transcription in individual live cells (Chubb et al., 2006; Golding et al., 2005) have shown that activation of almost all genes occurs in a pulsatile manner; a phenomenon known as transcriptional bursting. Use of the orthogonal MS2 and PP7 RNA tagging systems has demonstrated that numerous gene regulatory inputs are involved in producing a variety of bursting patterns (reviewed in Tunnacliffe and Chubb, 2020). Developmental changes in transcriptional bursting in live cells have previously been studied, mostly in *Drosophila* (Bothma et al., 2014; Lim et al., 2018; Scholes et al., 2019), *Dictyostelium* (Chubb et al., 2006; Muramoto et al., 2012; Tunnacliffe et al., 2018), and *Caenorhabditis* (Lee et al., 2019), while equivalent studies have not been reported in mammalian systems. To date, such studies have followed dynamic transcriptional changes occurring over relatively short time periods (1-2 h) or via sequential snapshots at different time points. This is largely due to the difficulty of conducting live imaging experiments over long time periods (up to days) without compromising cell viability or causing photobleaching. Consequently, studying transcriptional dynamics across the course of differentiation, with sufficient temporal resolution, is problematic, making the study of mammalian development particularly challenging.

Haematopoietic stem cells undergo lineage specification and, following commitment and differentiation, mature via a trajectory of well-defined morphological stages to produce ∼1-2 million red blood cells per second. This system is accurately recapitulated by various cell-based systems (Socolovsky et al., 2001, Zhang et al., 2003, Griffiths et al., 2012, Francis et al., 2020) and has established a general model for addressing how mammalian gene expression is regulated during changes in differentiation and development. Early in erythroid differentiation (hereafter referred to as erythropoiesis) a ∼65 kb topologically associating sub-domain (sub-TAD) containing the entire mouse *α*-globin locus forms (Brown et al., 2018; Hay et al., 2016). A cluster of 5 erythroid-specific enhancers are thereby brought into close physical proximity to the *α*-globin promoters to regulate their expression (Oudelaar et al., 2020). During the subsequent differentiation and maturation, including ∼4 cell divisions, globin gene transcription increases and eventually each erythroid cell accumulates up to ∼10,000 molecules of *α*-globin RNA (Orkin et al., 1975). However, the mechanisms by which different regulatory inputs associated with changes in transcriptional bursting control such gene expression in single cells is unknown.

By combining PP7 tagging of RNA transcripts and developing “on-microscope” cell staging, we were able to observe transcription dynamics of the mouse *α*-globin gene in real-time throughout sequential stages of erythropoiesis. We show that nascent α-globin transcription reaches a maximum between the early and intermediate stages of erythroid differentiation, preceding peak mRNA abundance and haemoglobin synthesis, and before transcription significantly declines at later stages of differentiation. The parameters used to define transcription in live cells are summarized in Figure 1. We find that changes in RNA production are primarily determined by the fraction of time a gene spends in an active transcriptional state (ON fraction), predominantly reflected in the burst frequency rather than the amplitude of the burst. Despite general trends in increased nascent transcription and ON fraction, we observed considerable variation in the patterns of transcriptional activity, as measured by the Fano factor (a measurement of variance/mean; Fig. 1), within and between cells at all stages of erythroid differentiation. Erythroid cells showed maximal transcriptional variability (also referred to as noise) both immediately before and after the peak in nascent transcription. Increased variability in these early and late cell stages is largely explained by the occurrence of sporadic, high amplitude bursts.

**Figure 1:**
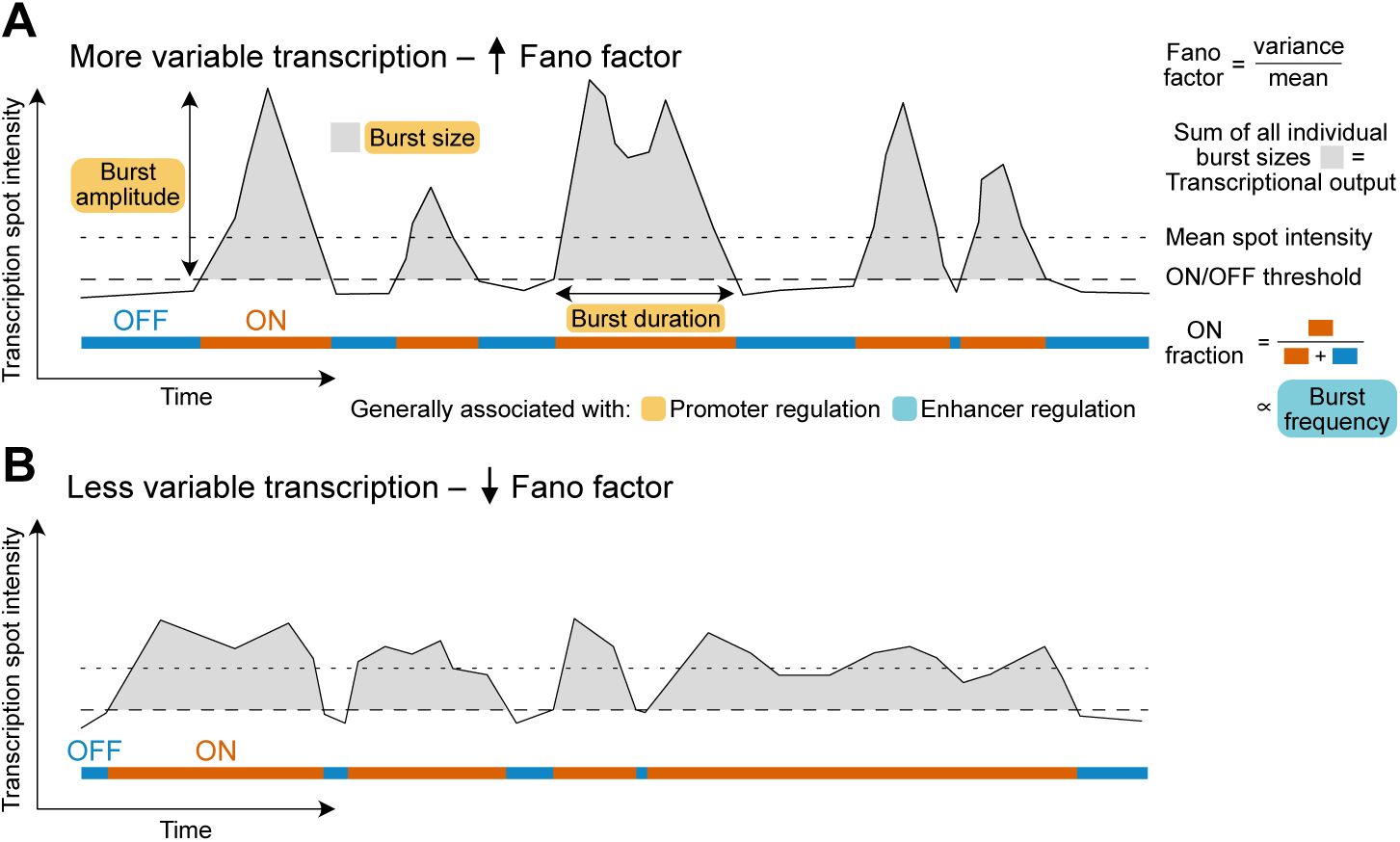
Schematic illustrating measurable features of transcription dynamics. Burst size changes are typically associated with promoter regulation, while burst frequency (ON fraction) is typically associated with enhancer regulation. More (A) and less (B) variable transcriptional activity traces of individual cells, with higher and lower Fano factor values respectively, are illustrated. For highly expressed genes, burst frequency can be difficult to measure directly from spot intensity traces. Measurement of ON fraction (similar to burst fraction in fixed cells, Bartman et al., 2016) in individual cells can be used to infer changes in burst frequency.

These findings suggest that the patterns of transcriptional bursting change during differentiation, and that variability in transcription is significantly reduced at the peak period of gene expression, perhaps via the establishment of a more stable interaction between enhancers and promoters within a transcriptional hub.

## Results

### Visualising α-globin transcription in live erythroid cells derived from mouse embryonic stem cells

To understand the kinetics of gene expression in individual live cells as they differentiate, we studied the α-globin genes (Fig. 2A), which are switched on and off at specific stages of erythropoiesis. This locus provides an extremely well characterized model that has established and illustrated many of the principles underlying mammalian gene expression (Oudelaar et al., 2021). To study α-globin transcription in real-time, we used the PP7 bacteriophage RNA labelling system (Lim and Peabody, 2002). We integrated a DNA sequence encoding an array of 24 PP7 loops into the first exon of a single allele of the *Hba-a1* gene in mouse embryonic stem (mES) cells using a recombinase-mediated cassette exchange (RMCE) strategy (Extended Data Fig. 1A) to generate *Hba-a1*-PP7 cells. Correct integration was confirmed via Southern blot and DNA FISH experiments (Extended Data Fig. 1B). To monitor changes in α-globin transcription throughout erythroid development, we used an *in vitro* mES cell differentiation system from which primitive erythroid cells are efficiently formed within embryoid bodies (EBs) during a differentiation period of up to 7 days (Francis et al., 2020; Keller et al., 1993; Keller, 1995). In this system, at a population level, both mature and nascent α-globin transcript levels increase significantly from day 4 to day 7, alongside well-characterised changes in expression of pluripotency and erythroid marker genes (Fig. 2B, Extended Data Fig. 2; Francis et al., 2020). Using this system, *Hba-a1-*PP7 mES cells differentiate normally along the erythroid lineage as assessed by immunophenotyping and morphology (Extended Data Fig. 1C, D). Furthermore, these cells exhibit a normal ratio of α/β-globin RNA expression (Fig. 2C) and chromatin accessibility of the locus (Fig. 2D). Therefore, labelling the endogenous *Hba-a1* α-globin gene using PP7 loops does not affect the activity of the locus or progression of erythropoiesis in differentiating mES cells.

**Figure 2:**
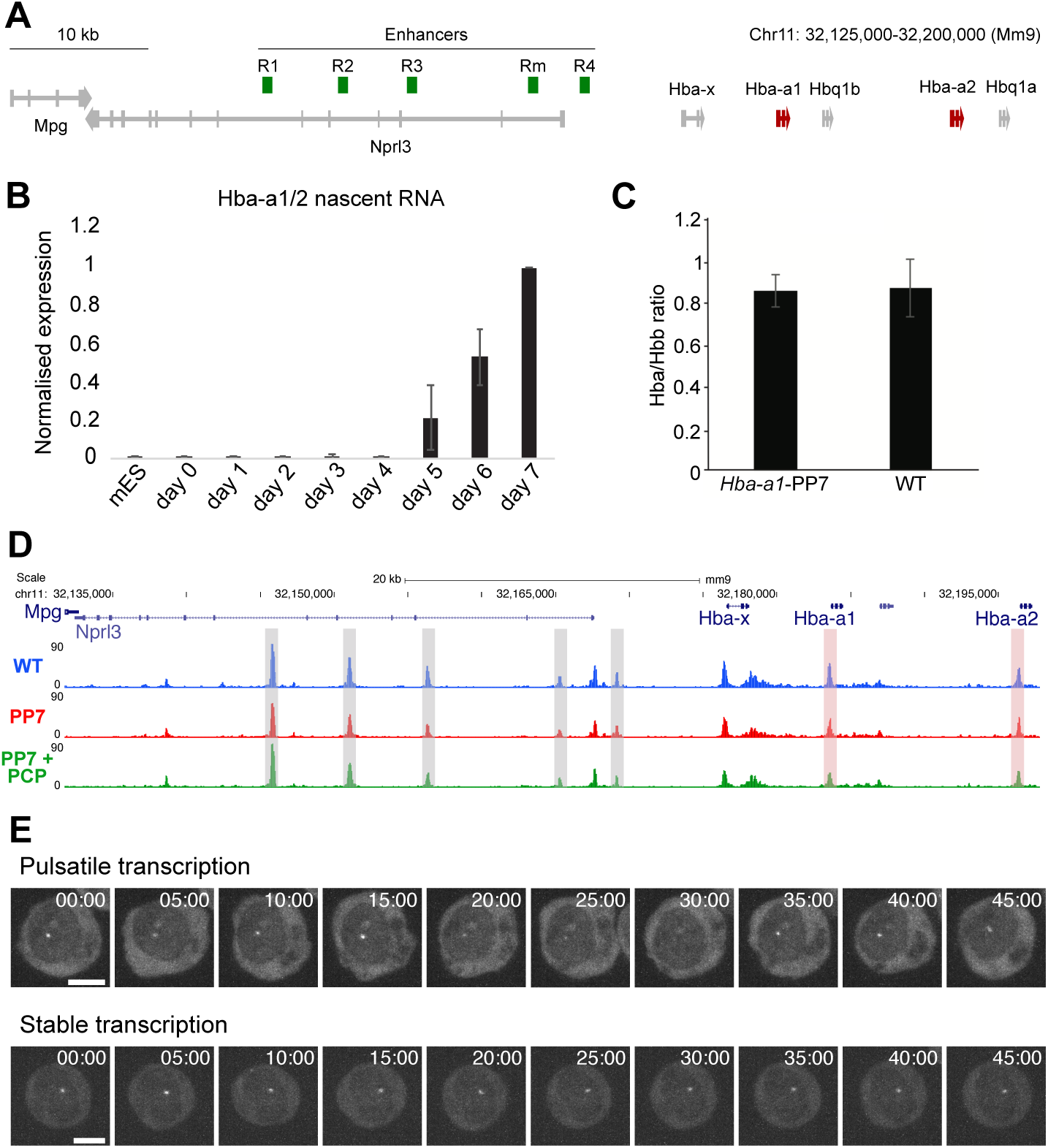
Visualising α-globin transcription dynamics in erythroid cells using the PP7 imaging system. (A) Schematic of mouse α-globin locus on chromosome 11. α-globin genes are highlighted in red, enhancers in green. (B) RT-qPCR time course of nascent α-globin mRNA during embryoid body (EB) differentiation. Data are normalised to Rn18s and then within differentiation time course. Data represent the mean of three independent experiments. Error bars = standard deviation. (C) RT-qPCR quantitation of α-globin mRNA abundance relative to β-globin. (D) ATAC-seq in WT, *Hba-a1*-PP7 (PP7) and *Hba-a1*-PP7 + PCP-GFP (PP7 + PCP) cells at the α-globin locus. Grey shaded regions show known enhancers, red shaded regions show α-globin genes. (E) Representative examples of nascent *Hba-a1*-PP7 transcription over time showing pulsatile or more stable gene activity in individual erythroblasts derived from day 6 EBs. Frames are a maximum projection of five z-slices around transcription spot from within the full image stack. Time in minutes. Scale = 5 μm.

To detect *Hba-a1* transcription in the *Hba-a1-*PP7 clone, a transgene encoding a constitutively expressed PP7 coat protein fused to GFP (PCP-GFP) was randomly integrated into the genomes of both wild-type (WT) and *Hba-a1-*PP7 mES cells. Clones exhibiting medium levels of GFP expression and relatively uniform expression across the population were chosen for analysis. These *Hba-a1-*PP7 + PP7 coat protein (PCP)-GFP cells differentiate normally to EBs (Fig. 2D, Extended Data Fig. 1C, E). We confirmed that a nuclear-localised transcription spot was only detected in differentiated cells containing PP7 loops integrated in *Hba-a1*, and not in WT control cells (Extended Data Fig. 3A). These transcriptional foci remain localised within the nucleus throughout the time course of the experiment (Extended Data Fig. 3B, C). However, as also observed by others in yeast (Heinrich et al., 2017; Tutucci et al., 2018), some cells exhibited cytoplasmic fluorescent foci. Such foci were excluded from further analysis (see Methods). Initial time-lapse experiments showed that the pattern of *Hba-a1* activity within individual cells derived from day 6 EBs is variable. In the course of one hour, some cells displayed pulsatile gene activity, and others showed a more stable, albeit still variable, pattern of transcription (Fig. 2E). This showed that *α*-globin may exhibit a range of transcriptional bursting behaviours in erythropoiesis.

### On-microscope staging of erythropoiesis

While numerous studies have investigated the dynamics of gene transcription in mammalian cell culture (Fritzsch et al., 2018, Harper et al., 2011, Molina et al., 2013, Ochiai et al., 2014, Rodriguez et al., 2019, Suter et al., 2011, Zoller et al., 2015), live-cell studies of transcription dynamics during mammalian differentiation have not been reported. To study the kinetics of α-globin transcription at different stages of erythropoiesis, we initially imaged transcription within individual cells derived from day 5, 6 and 7 EBs for 1 hour with 5 minute (min) frame intervals. We observed considerable variability in the dynamics of *α*-globin transcription between live cells when simply stratifying by days in culture (Extended Data Fig. 4). We hypothesised this variability was most likely due to the presence of a mixture of cells at different stages of differentiation at each time point as a result of unsynchronized differentiation within EBs.

This problem is commonly encountered in live imaging studies of dynamic cell processes and so, to overcome this, we stratified erythroblasts obtained from these cultures using directly-conjugated antibodies (Eilken et al., 2009) that recognize CD71 and Ter119, known surface markers of erythropoiesis whose levels change in a predictable manner during erythroid differentiation (Socolovsky et al., 2001; Zhang et al., 2003, Fraser et al., 2006, Chao et al., 2015). Although erythroid cells can be separated in various stages of differentiation using fluorescently activated cell sorting (FACS), we observed reduced viability of sorted cells in prolonged imaging experiments, potentially caused by an extended period of sorting-associated stress. Therefore, we developed live-cell antibody staining to stage cells throughout erythropoiesis directly under the microscope (see Supplementary Note for further discussion).

We assessed the ability of fluorescence microscopy to enable accurate cell staging during erythropoiesis. Measuring the levels of CD71 and Ter119 in a single 3D image stack (Fig. 3A) from FACS sorted populations (F1-F6) enabled erythroid-cell staging similar to that obtained using FACS (Fig. 3B; Extended Data Fig. 5A, B), giving access to a full spectrum of erythroid differentiation states. May-Grünwald-Giemsa (MGG) staining of the same FACS-sorted populations (F1-F6) showed CD71^low^/Ter119^-^ cells (F1) were largely comprised of proerythroblasts, while CD71^high^/Ter119^high^ cells (F6) represented later erythroblast (poly/orthochromatic erythroblast) stages, with a smooth progression in the changing proportion of cells within intermediate populations (Fig. 3C, D and Extended Data Fig. 5C, D, E). Staining live cells by directly-conjugated antibodies targeting CD71 and Ter119 therefore enabled rapid staging of EB-derived erythroid cells into early, intermediate and late stages of differentiation by fluorescence microscopy.

**Figure 3:**
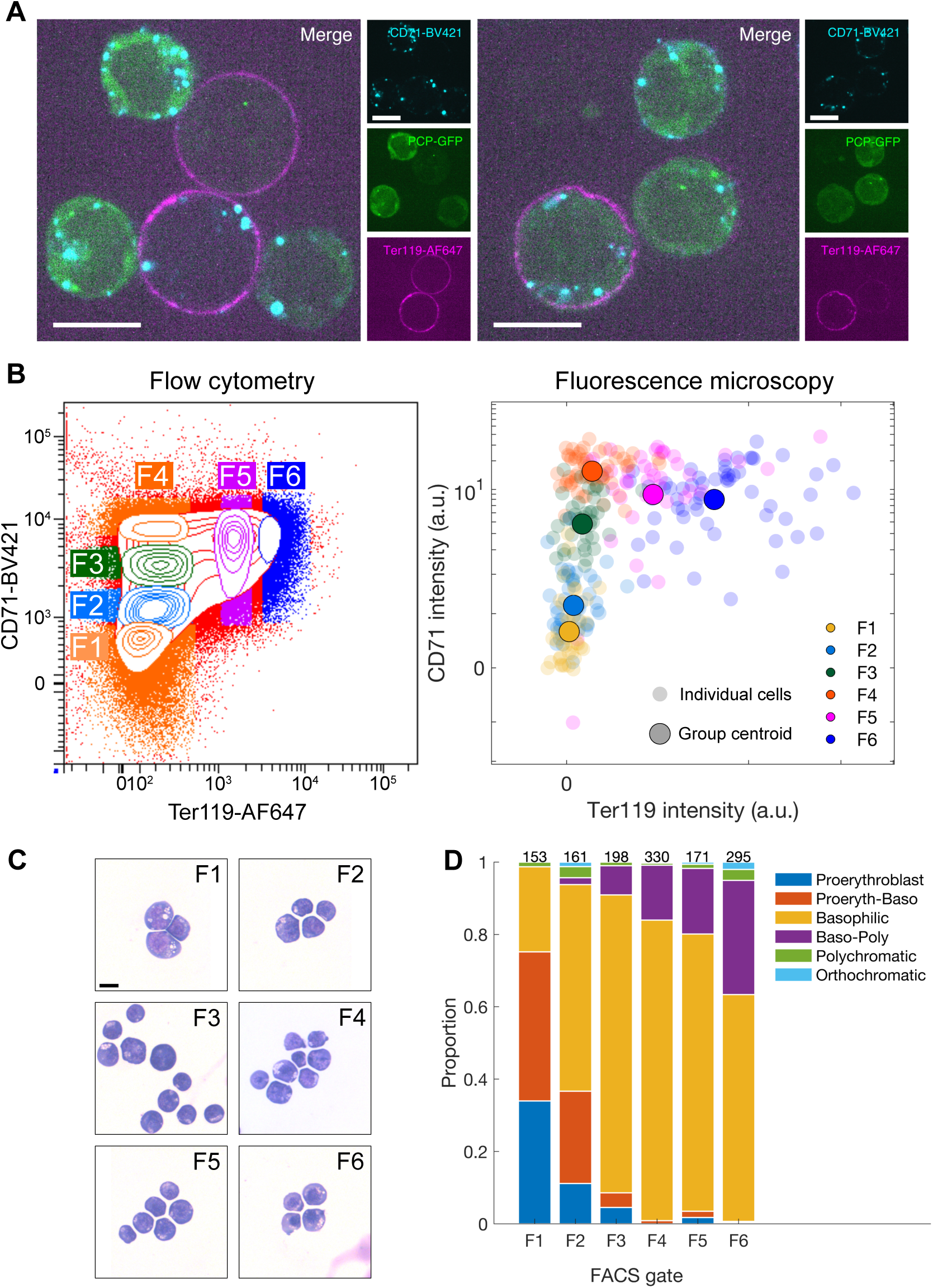
Live-cell antibody staining enables on-microscope staging of erythroid cells. (A) Example images of cells with variable levels of CD71 and Ter119 staining. Single z-slices shown. Scale = 10 μm. (B) Day 7 EB-derived cells FACS-sorted (F1, F2, etc) according to CD71/Ter119 staining (left) and subsequently imaged on the microscope (right). (C) Representative images of cells from the sorted populations in B stained with May-Grünwald-Giemsa solution. Scale = 10 μm. (D) Proportion of erythroblast stages in each FACS-sorted population scored by morphology from MGG-stained cells in C. Numbers indicate number of cells scored.

### Stratifying cells into progressive stages of differentiation

Having shown that live-cell antibody staining enabled on-microscope identification of erythroblast stages, we next wanted to simultaneously monitor nascent transcription and the stage of differentiation of individual erythroid cells. We imaged *Hba-a1*-PP7 transcription for 1 hour with 2.5 min frame intervals and subsequently collected single stacks of CD71 and Ter119 markers at day 6 of EB differentiation (Extended Data Movies 1-4). The time frame interval of 2.5 min was optimized to capture the majority of transcriptional bursts while minimising photobleaching (Extended Data Fig. 6).

To enable stratification of these cells into progressive differentiation stages, we plotted cells onto a CD71/Ter119 axis (Extended Data Fig. 7A). We then mapped each cell onto a one-dimensional differentiation axis from the two-dimensional CD71 and Ter119 staining pattern. To stage the cells, we used an empirically defined series of curves that follow the changes in intensity of CD71 and Ter119 markers through erythroid differentiation (Extended Data Fig. 7A; Francis et al., 2020). Multiple curves were employed to accommodate the broad distribution of CD71 staining in Ter119^low^ cells. The differentiation stage of each cell was estimated from its location with respect to the nearest curve (Extended Data Fig. 7B, C). Cells were then grouped into 6 stages of erythroid differentiation (DS1-DS6) according to their positions along this differentiation axis (Extended Data Fig. 7D, E). This showed a peak in the number of cells at intermediate stages of erythroid development (Extended Data Fig. 7D) consistent with previous studies of EBs at Day 6 of differentiation (Francis et al., 2020).

To validate this cell stratification approach, we compared cells at DS1-DS6 with conventionally defined erythroid precursors. We mapped cells from the FACS-sorted populations (F1-F6) imaged by microscopy onto a differentiation axis in the same way as for DS1-DS6 (Extended Data Fig. 7F) and aligned the two axes (see Methods) to allow direct comparison between the two datasets. Subsequent superimposition of the relative proportions of erythroblast stages from F1-F6 populations (Fig. 3C, D) onto histograms of their position in differentiation (Extended Data Fig. 7Gi, ii) enabled us to approximate the differentiation stage of cells in DS1-DS6 (Extended Data Fig. 7Giii). Using these approaches, we established that DS1 represents proerythroblasts, cells at stage 3-5 (DS3-DS5) broadly correspond to basophilic erythroblasts, and those at stage 2 (DS2) represent the transition between these two. Stage 6 (DS6) represents a mixture of later erythroblast stages including poly/orthochromatic erythroblasts and some basophilic erythroblasts (Extended Data Fig. 7Giii). Although not providing a perfect separation, our on-microscope staining approach enabled live imaging of transcriptional dynamics of cells at all stages of terminal erythropoiesis.

### Variable patterns of transcription within individual cells are seen throughout erythropoiesis

Stratifying individual cells into differentiation stages (DS1-DS6) using on-microscope analysis (Extended Data Fig. 7D, E) enabled us to test in further detail how *α*-globin transcription changes at specific points in erythropoiesis. For each cell analysed, we determined the average spot intensity and the fraction of time spent in transcription (ON fraction) after setting an ON/OFF threshold. For individual bursts, we recorded the duration and size (Fig. 1). Importantly, data were analysed using multiple ON/OFF thresholds to ensure robust results. In general, we found that cells at the mid stages of erythropoiesis (DS3 and DS4) exhibit the highest transcription spot intensity (DS1 vs. DS4 median of mean spot intensities: 478.9 +/-506.9 vs 670.7 +/-626.7 respectively) (Fig. 4A, Extended Data Fig. 8A, B) and more frequently reach higher burst amplitude (a visual estimation of brighter colours in binned data in Fig. 4B, and raw data Extended Data Fig. 8C) in individual cells. On average, cells at stage DS3 and DS4 are active 72-73% of the time (high ON fraction), compared to only 43% and 33% for stages DS1 and DS6, respectively (Fig. 4C, Extended Data Fig. 8D, E). These results are consistent with experiments in fixed cells where, at any one time, 70-80% of cells were found to be transcribing globin genes at the peak of activation (Bartman et al., 2016; Brown et al., 2006). Together, these findings match the changes in mean transcription levels (Fig. 4A) and ∼90% of the variability in mean spot intensity was explained by the fraction of time spent in the ON state (Fig. 1, Extended Data Fig. 8Ei). It has previously been suggested, based on smFISH analysis, that the fraction of cells transcribing at any one time (called burst fraction Bartman et al., 2016) is related to burst frequency. Our use of the ON fraction is essentially a measurement of the burst fraction in live cells. While the fraction of time a gene spends active could also be affected by the duration of bursts, measurement of individual bursts showed that this parameter is largely invariant across differentiation (see below). Therefore, our data suggest that regulation of burst frequency is the primary determinant of the average levels of α-globin nascent transcription during erythropoiesis.

Having shown that average levels of nascent α-globin transcription (mean spot intensity) change throughout differentiation, we next wanted to determine whether the dynamic patterns of transcription within individual cells also change throughout differentiation. Fluctuations in transcriptional activity in individual cells, or “transcriptional variability”, can be measured by the Fano factor (variance divided by the mean, σ/μ) which is a measure of noise-to-signal ratio (see example traces Extended Data Fig. 8F). Using this metric we were able to assess intra-cellular variability in transcriptional activity of the gene during the imaging period (Fig. 1). In general, we found that at each stage of erythropoiesis, as gene expression increases, so does the Fano factor (Extended Data Fig. 8Fi). However, we found marked differences in the relationship between the Fano factor and the level of transcription at different times in erythropoiesis. At the peak of transcriptional activity in the population, the gradient of the linear regression between transcription (mean spot intensity) and the Fano factor within individual cells was around half that seen at the differentiation stages flanking the peak (stage 2 = 0.54 vs. stage 3 = 0.28; Fig. 4D, E, Extended Data Fig. 8G). Therefore, in the early stages of erythropoiesis (DS1 and DS2), when the α-globin genes first become transcriptionally active, the noise-to-signal ratio (Fano factor) is high relative to that which would be predicted from the level of expression (Extended Data Fig. 8F). Transcription then becomes less variable when the genes are fully active (DS3 and DS4), and again becomes increasingly variable as the genes are being switched off (DS5 and DS6) (Fig. 4D, E). These changes in the patterns of transcription, consistent with previous RNA FISH studies, suggest differences in the molecular mechanisms of α-globin activation in single cells at each stage as they progress through erythropoiesis.

**Figure 4:**
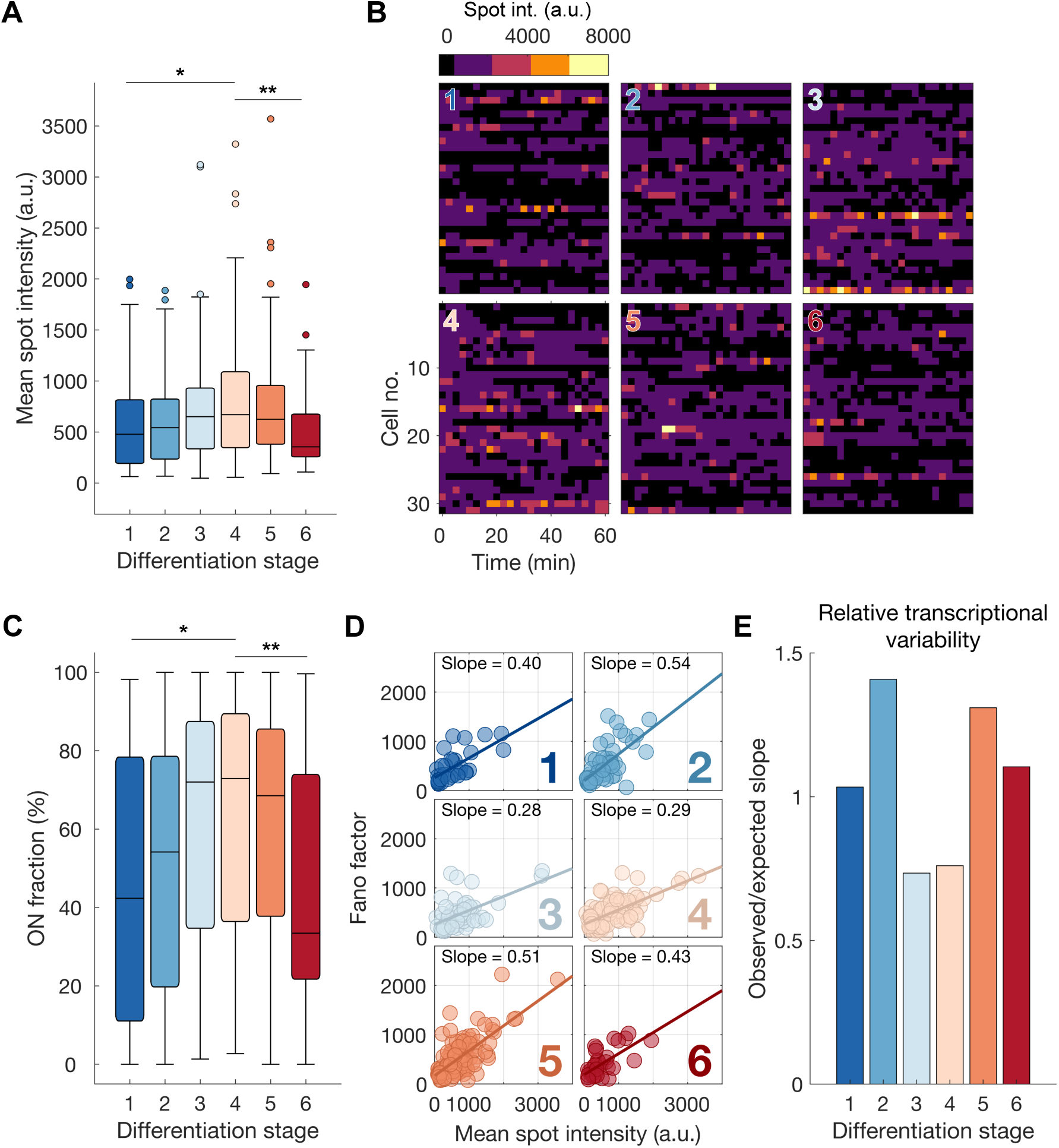
Modulation of α-globin transcription dynamics throughout erythropoiesis. (A) Distribution of mean spot intensities of individual cells through different stages of erythropoiesis. a.u. = arbitrary units. (B) Example spot intensity traces of cells within differentiation stages. Rows represent individual cells. Intensities are binned above the ON threshold of 350 arbitrary intensity units (black). (C) Distribution of the proportion of time spent in the active ON transcriptional state (above ON threshold) for each cell by differentiation stage. (D) Relationship between mean spot intensity and intra-cellular transcriptional variability (Fano factor) for individual cells across differentiation stages. (E) Relative levels of transcriptional variability throughout erythropoiesis defined by the ratio of the observed slopes of linear regression in D compared to that of all cells pooled (expected). Boxes within boxplots show median and interquartile range, whiskers show 9^th^ and 91^st^ percentile of distribution. Statistical comparisons are two-sided Mann-Whitney U tests where * p < 0.05, ** p < 0.01.

### Characterising transcriptional bursting in individual cells

To investigate potential mechanistic differences in transcription at each of these stages, we analysed the relationship between different parameters of α-globin transcription dynamics (i.e. burst size and frequency, see Fig. 1) within individual cells during erythropoiesis. Regulation of these dynamics has generally been linked to the action of transcription factors (TFs) at promoters and enhancers (Fig. 1). Burst size, which can be broken down further into burst duration and burst amplitude, is thought to be regulated via the promoter. By contrast, burst frequency is thought to be determined by enhancers (Fig. 1) (Bartman et al., 2016; Donovan et al., 2019; Fukaya et al., 2016; Stavreva et al., 2019).

We asked whether changes in burst frequency were responsible for the differences in variability during differentiation. We have shown that, throughout differentiation, the ON fraction (which is related to burst frequency, Bartman et al., 2016), is strongly correlated with mean gene activity measured from the transcription spot intensity (Fig. 4A-C, Extended Data Fig. 8Ei) suggesting that increasing burst frequency explains overall levels of *α*-globin RNA synthesis as erythropoiesis proceeds. By contrast, the ON fraction shows only a very weak correlation with transcriptional variability (Extended Data Fig. 8Eii) suggesting that burst frequency does not account for the observed differences in variability at different stages of differentiation.

To examine changes in burst size during erythropoiesis, we first measured the total transcriptional output (area under the curve between spot intensity fluctuations and above the ON threshold, Fig. 1) relative to the total ON duration in individual cells over a period of 60 minutes (Fig. 5A, B). This enabled us to assess differences in the average burst amplitude for each cell, since total transcriptional output is a product of the duration and amplitude. We estimated the average relationship between duration and amplitude across all cells using local regression and observed a clear inflection point in the gradient of this relationship (Fig. 5A). A higher gradient means increased transcriptional output for a given change in duration, and therefore indicates a higher average burst amplitude. Cells lying to the right of the inflection point which are almost continuously active (which we call ‘high ON’ cells, active >89% of the time see Supplementary Note), therefore have an increased burst size as a consequence of increased burst amplitude. By contrast, ‘low ON’ cells to the left of the inflection point (Fig. 5A; active <89% of the time) have much lower average burst amplitude. In summary, these findings show that burst frequency is not primarily responsible for the observed differences in variability at different stages of differentiation. However, burst amplitude of α-globin transcription appears to vary with the duration of time for which the gene is active (Fig. 5A). This prompted us to examine if the burst amplitude may explain the different degrees of variability that we observe at different stages of erythropoiesis.

**Figure 5:**
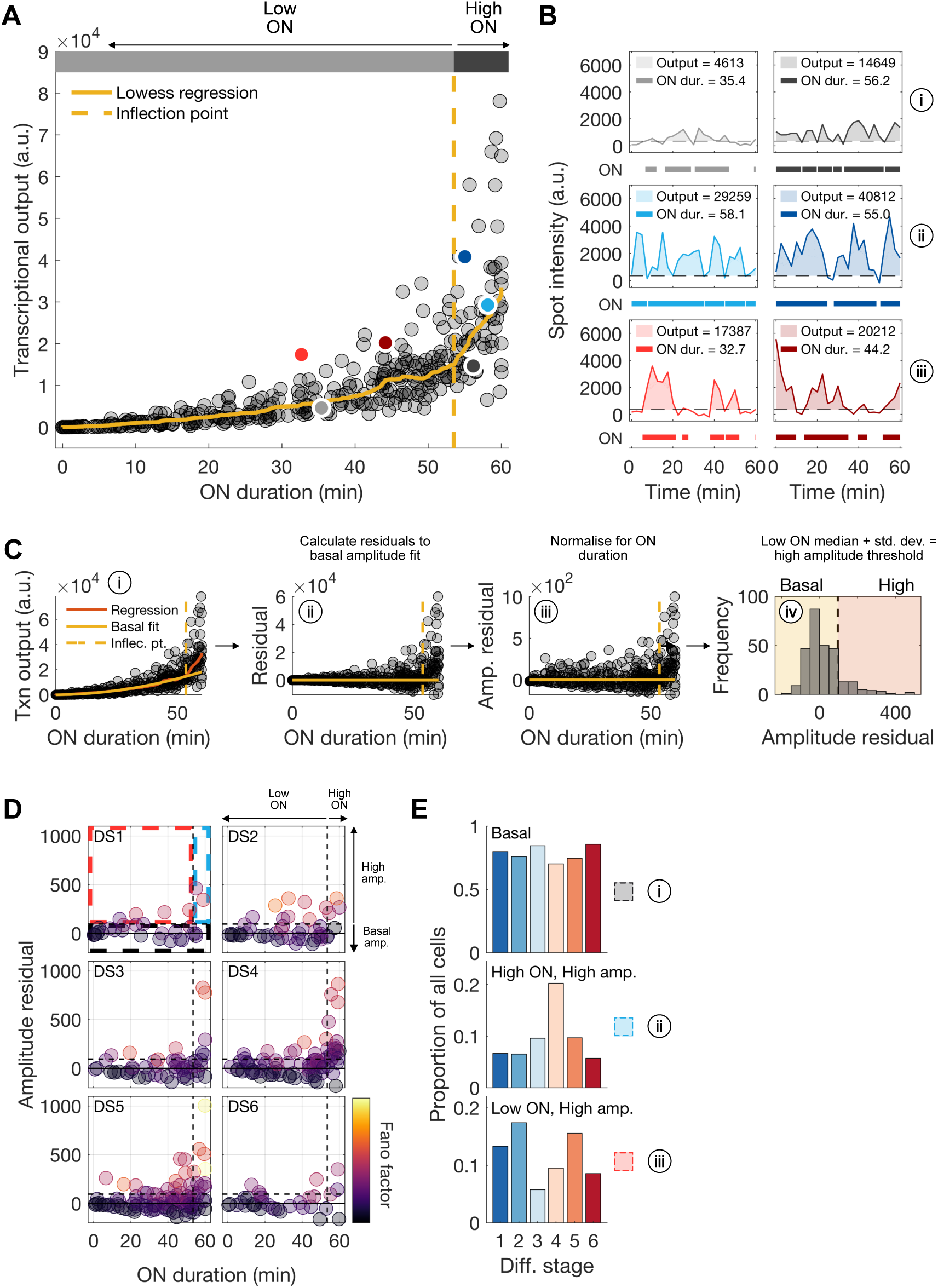
Changes in transcriptional behaviour during erythropoiesis. (A) Relationship between ON duration and total transcriptional output (area below the curve and above ON threshold for entire imaging period for each cell). LOWESS regression outlines the local relationship between these variables. The inflection point marks a step-change in the gradient of the regression. Cells above the inflection point we call ‘High ON’ cells, while those below are ‘Low ON’. a.u. = arbitrary units. (B) Example spot intensity traces for individual cells marked by coloured circles in A. Below each panel, thick lines indicate when intensity traces for each cell are above ON threshold (dotted lines). The measured ON duration and area under the curve (shaded area) are given for each example cell. (C) Method for calculation of the high amplitude threshold, above which cells are said to have high average burst amplitude. The relationship between ON duration and transcriptional output for basal amplitude transcription was estimated by fitting a quadratic curve to local regression data points in ‘Low ON’ cells (yellow solid line, i). For each cell, residuals of transcriptional output to this basal amplitude estimate were calculated (ii) and normalised for ON duration (iii). This ‘amplitude residual’ represents the deviation of each cell from a basal amplitude bursting behaviour, i.e. if the amplitude residual is close to 0 then a cell is exhibiting basal amplitude bursting. ‘High amplitude’ cells are those with an amplitude residual of greater than one standard deviation above the median (iv). (D). Amplitude residuals as a function of ON duration for cells in each differentiation stage. Data points are coloured by the transcriptional variability (Fano factor) for each cell. Coloured dashed lines indicate basal amplitude cells (black), low ON high amplitude cells (red), high ON high amplitude cells (blue), demarcated by the previously defined low/high ON and basal/high amplitude thresholds. (E) Calculated proportions of cells with different bursting behaviours at each differentiation stage as defined by dashed regions in D.

### Coincident changes in burst amplitude and variability during erythropoiesis

To further analyse the role of burst amplitude in generating variability in transcription throughout erythropoiesis, we examined transcriptional burst traces in individual cells. While in most ‘low ON’ cells, expression fluctuates around a basal level of transcription (Fig. 5Bi), higher amplitude bursts were observed not only in ‘high ON’ (Fig. 5Bii) cells but also in a proportion of ‘low ON’ cells (Fig. 5Biii). We used the distinction between ‘high ON’ and ‘low ON’ cells (Fig. 5A) to quantify the proportion of cells exhibiting these different transcriptional behaviours across the differentiation stages (Fig. 5C and Supplementary Note). Most cells (>70%) display uniformly low burst amplitude throughout the imaging period (“basal”, Fig. 5Bi, D and Ei). Consequently, such cells have a low transcriptional noise to signal ratio (Fano factor; Fig. 5D, Extended Data Fig. 9A). The majority of cells at all stages of differentiation fall into this category (Fig. 5D and E). A smaller proportion of cells (6-20%) show near-continuous activity with regular, intense bursts of transcriptional activity leading to increased transcriptional output (Fig. 5Bii, D and Eii). We define these cells as ‘high ON, high amplitude’ cells. Although this is associated with an increase in the Fano factor this is *relatively* low compared to the level of expression (Fig. 4E, 5D). This pattern is most prominent at stages DS3-4 of differentiation when *α*-globin transcription reaches maximal levels (Fig. 5D, E). In the remaining cells we see sporadic, strong bursts of transcription separated by periods of transcriptional quiescence (‘low ON, high amplitude’ cells; Fig 5Biii, D and Eiii). These are most prominent at stages DS1 and 2 (13-17% of cells), when transcription is starting, and also at stage DS5 (16% of cells) as the genes are being downregulated (Fig. 5D, E). At these time points, variability is *relatively* high compared to transcriptional output (Fig 4E, 5D). A summary of these data is presented in Extended Data Figure 9D. We observed that the proportion of these low ON cells exhibiting high amplitude bursts seemed to match the trends in transcriptional variability across differentiation (Fig. 4E, 5E).

To test whether burst amplitude is indeed higher at stages DS2 and 5, when transcription is more sporadic, we characterised individual bursts in cells at all differentiation stages (Extended Data Fig. 10A). The duration of individual bursts was largely invariant across the differentiation stages (median 4.3-5.6 min) suggesting that this parameter of burst control is unlikely to be extensively regulated during erythropoiesis (Extended Data Fig. 10C). In general, high ON cells have a higher burst size for a given burst duration (indicating a higher burst amplitude) than low ON cells, in keeping with our earlier analysis (Extended Data Fig. 10D, Fig. 5A). Furthermore, the relative burst amplitude for low ON cells is highest in differentiation stages DS2 and DS5, matching the trends in noise during differentiation (Extended Data Fig. 10E, F, Fig. 4E, 5E).

Together these data suggest that increased burst amplitude in ‘low ON’ cells (those exhibiting sporadic transcriptional bursts) could be responsible for changes in transcriptional variability during erythropoiesis. Most importantly, the time spent in an active transcriptional state (ON fraction), most probably linked to enhancer-promoter communication, appears to be the dominant control point for α-globin transcription levels during differentiation.

## Discussion

We have studied the real-time dynamics of transcription during the process of differentiation in individual living cells. In contrast to previous studies, we have been able to determine the pattern of nascent transcription that occurs over several days as mammalian stem cells progress to fully differentiated mature cells accumulating large amounts of mRNA and protein. The key to this was our ability to accurately define erythroblast stages from a heterogeneous population under the microscope using a combination of antibodies to two well-characterised cell surface markers (CD71 and Ter119). This enabled us to define the differentiation stage of individual cells and correlate this with previously established morphological criteria. This in turn allowed us to develop an approach that facilitated subsequent analysis of live imaging of transcription dynamics throughout erythropoiesis.

The *in vitro* erythroid body culture system used here recapitulates primitive erythropoiesis (Francis et al., 2020), which is broadly similar to definitive erythropoiesis in terms of its core regulatory modules (Kingsley et al., 2013) and temporal pattern of CD71/Ter119 expression (Pijuan-Sala et al., 2019). Although we have shown that changing levels of CD71 and Ter119 enable visualisation of maturing erythroblast stages in our system, this is still a relatively coarse dissection of erythropoiesis. However, we have shown that the use of live-cell antibody staining for staging cells directly under the microscope facilitates live imaging studies over a prolonged period of differentiation. Given the simplicity of the method (Eilken et al., 2009), this approach will greatly extend and improve the application of single-cell transcriptional dynamics in general (see Supplementary Note for further discussion).

Studying *α*-globin expression at well characterized stages of erythropoiesis we found that, consistent with fixed cell imaging, nascent transcription (as measured by spot intensity) reaches a maximum at the intermediate stages of erythropoiesis. We also found that the main factor driving the level of α-globin transcription in any cell is the fraction of time spent in the ON state, given that the peak in both average transcription and ON fraction occurred in the mid stages of differentiation, with the average likelihood of a cell transcribing at this stage being ∼70% (Figure 4, Extended Data Figure 8). This pattern of α-globin expression is consistent with previous fixed-cell RNA FISH experiments describing transcription dynamics at several time points during blood differentiation where 70-80% of cells were found to be transcribing both α-globin and β-globin at the peak of activity (Bartman et al., 2016, Brown et al. 2006).

The patterns of nascent transcription observed here in real-time, were not as different at each stage of differentiation as might have been expected. Further stratification of erythroid differentiation would be expected to reveal even greater differences in patterns of nascent transcription. Each stage of differentiation included cells with three broad categories of transcriptional bursting (Fig. 5B). In most cells (>70%) at all stages of differentiation, we saw low levels of transcriptional bursting (basal Fig. 5Bi). In a small proportion of cells (6-20%) we saw higher average levels of expression punctuated with intense bursts of transcriptional activity (Fig. 5Bii, high ON duration, high amplitude bursting). This is most marked at the intermediate stage of differentiation (DS4) when spot intensity and the ON fraction is at its maximum. Finally, in the remaining cells we see sporadic bursts of transcription separated by periods of transcriptional quiescence (Fig. 5Biii): these are most prominent at stages DS1 and DS2 (13-17% of cells), when transcription is starting, and also at stage DS5 (16% of cells) as the genes are becoming inactive. This suggests that between individual cells at similar stages of differentiation there is a considerable amount of variation in the patterns of transcription. The data presented here suggest that in contrast to analyses of cell populations, which suggest transcription and the accumulation of RNA progresses in a uniform manner, the underlying process of transcription is inherently variable. Ultimately, such variability in transcription may be mitigated by post-transcriptional mechanisms, not least by changes in the relative stability of different mRNAs, leading to the accumulation of globin RNA during erythropoiesis (Waggoner and Liebhaber, 2003).

The different patterns of transcriptional bursting observed here in real time, as opposed to previous studies of fixed cells, suggest that the molecular mechanisms underpinning α-globin transcriptional activation may change during erythropoiesis. Detailed understanding of the mechanisms regulating globin gene expression throughout erythropoiesis enable us to consider how these may explain the changing dynamics of *α*-globin transcription. In general, enhancers have been shown to primarily mediate changes in the frequency and probability of transcriptional bursts (Bartman et al., 2016; Fukaya et al., 2016; Rodriguez et al., 2019). Enhancers most frequently influence transcription by coming into close proximity to their cognate promoters (Furlong and Levine, 2018), although exactly how this relates to transcriptional bursting is uncertain (Alexander et al., 2019; Chen et al., 2018). During erythropoiesis, the α-globin enhancers come into increasingly frequent proximity to the α-globin promoters (Oudelaar et al., 2020) within a self-interacting domain (Brown et al., 2018). The frequency of proximity increases in line with increased globin transcription. We have shown here that burst frequency appears to be the primary control point for α-globin transcription, suggesting that activation of the promoters by the enhancers increases the ON duration in early erythropoiesis.

It has previously been suggested that, for cells to increase transcription from a gene such as α-globin, burst frequency is first increased until a threshold is reached, above which burst size is preferentially increased (Dar et al 2012). Our data agree with this idea (Figure 5), with ‘low ON’ cells more likely to transcribe with basal amplitude while ‘high ON’ cells are more likely to exhibit high amplitude bursting. This suggests that, in the case of α-globin, the threshold proposed by Dar et al. (2012) occurs when the gene is almost continuously active, beyond which burst frequency cannot be increased further. This switch from a lower to a higher burst amplitude in high ON cells suggests α-globin can be transcribed at multiple different initiation rates as measured by the PP7 system. This phenomenon has been demonstrated for several other genes (Tunnacliffe and Chubb, 2020).

The decrease in α-globin ON duration at stages 5 and 6, could be due to mechanisms associated with chromatin condensation as cells mature into late erythroblast stages (Mei et al., 2021). The mechanism behind the increase in likelihood of infrequent high amplitude bursts and transcriptional variability when the genes are not continuously active, either side of the maximum α-globin activity in the middle of terminal erythropoiesis is less clear. One component of this might be mediated via chromatin accessibility to TFs increasing as erythropoiesis progresses and then decreasing as the erythroid nucleus condenses. Recent imaging studies have highlighted TF dwell time as a key determinant of burst size in both yeast and mammalian cells with more stable binding leading to increased output (Donovan et al., 2019; Stavreva et al., 2019). Furthermore, in keeping with our study, a single inducible gene was shown to exhibit a range of noise-mean (Fano factor) relationships in transcription, depending on the concentration of a particular TF (Hansen and Zechner, 2021). Recent work also suggests that the formation of transcriptional compartments, containing a high concentration of TFs may increase the efficiency of transcription (Boija et al., 2018, Garcia et al., 2021). Our observations are therefore consistent with transient formation of a TF-concentrating transcriptional compartment initially giving rise to variable transcription followed by the formation of a more stable compartment producing higher levels of transcription with less transcriptional noise (Figure 6). Transcription again becomes variable as the gene expression decreases towards the end of differentiation.

**Figure 6:**
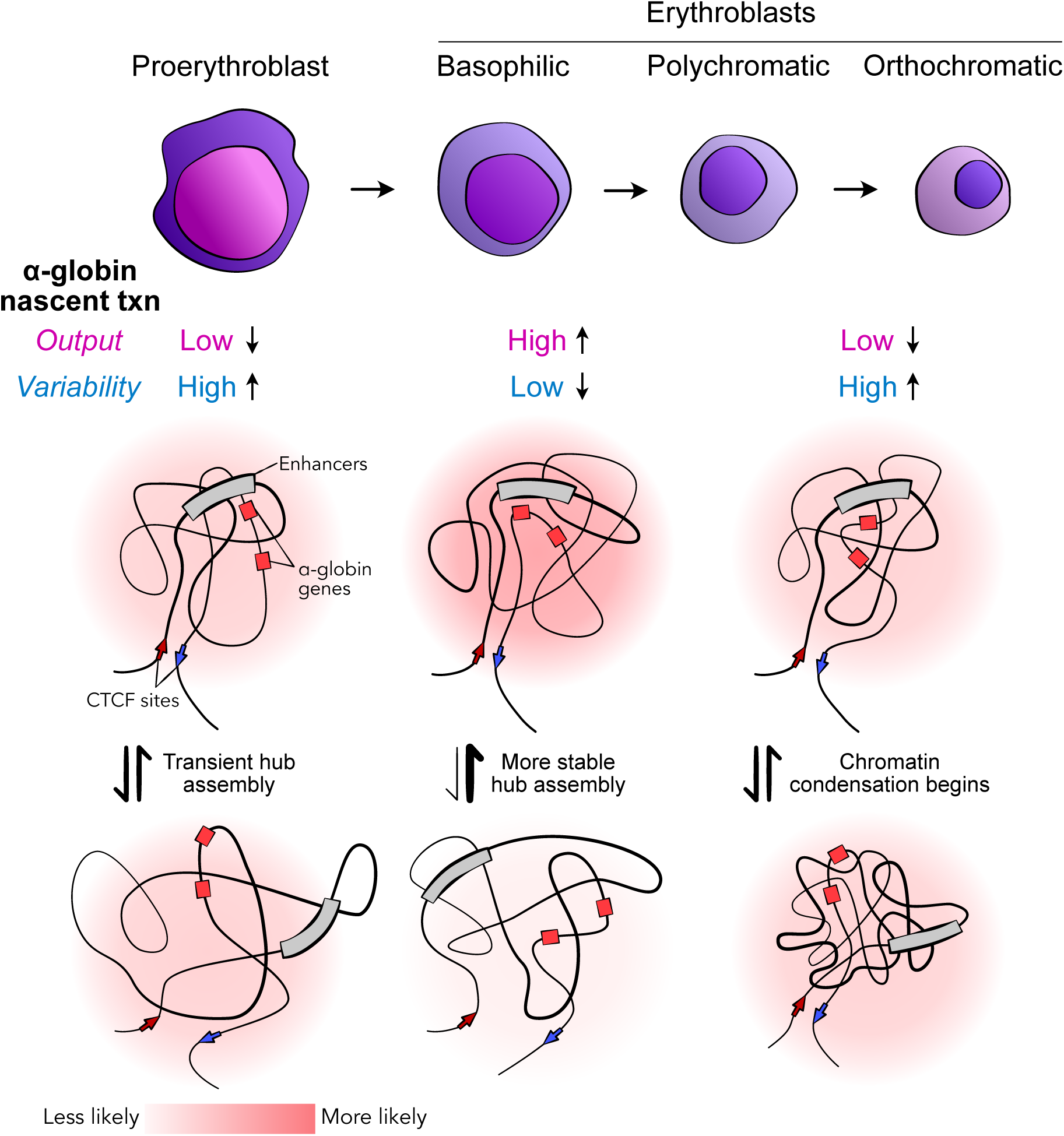
Schematic of proposed model for the observed changes in transcriptional output and variability as cells progress through erythropoiesis. Initially, assembly of a transcriptional hub occurs transiently causing an increase in intra-cellular transcriptional variability (left). More stable hub assembly occurs in the mid stages of differentiation in conjunction with an increased transcriptional output and reduced levels of relative intra-cellular variability (middle). Towards the latter stages of differentiation, chromatin condensation begins and the transcriptional hub is less stable, again causing an increase in relative intra-cellular transcriptional variability.

In summary, this study introduces technical advances for imaging of transcription dynamics during differentiation and development and provides fundamental insights into how dynamic patterns of transcription of a developmentally regulated gene are altered during differentiation.

## Methods

### Constructs

The targeting construct to establish the acceptor site for RMCE at the α-globin locus was generated in sequential steps using λ-red-mediated recombineering. First, a Tn10-rpsL-gentR cassette flanked by AscI sites was inserted in place of the *Hba-a1* gene sequence (coordinates 32,182,681-32,185,338 in mouse reference genome, build mm9) in a mouse RP22-289A22 BAC clone (BACPAC Resources Centre). Subsequently, the integrated cassette and flanking 4 and 6.9 kilobase (kb) blocks of adjacent BAC-derived *Hba-a1* sequence was retrieved by gap repair recombineering into a p15A minimal vector. Finally, the Tn10-rpsL-gentR cassette was replaced in the p15 plasmid with a pgk-Hyg-TK cassette that contains a PGK promoter and linked sequence encoding a fused hygromycin phosphotransferase and HSV thymidine kinase protein flanked by loxP and lox511 sites, amplified from a modified ZRMCE vector (a kind gift from Ann Dean). Individual recombineering and cloning steps were assessed by antibiotic selection, PCR amplification, restriction enzyme digestion, and sequencing. Thus, the final p15A-A22BAC-loxP-pgk-HygTk-lox511 construct was generated with 4 and 6.9 kb homology arms for gene targeting into the *Hba-a1* locus in order to make the RMCE acceptor site.

The *Hba-a1*-PP7 construct to be used as the donor vector for RMCE was generated in sequential steps using λ-red-mediated recombineering. First the region of *Hba-a1* gene sequence was retrieved from BAC clone RP22-289A22 equivalent to that sequence replaced above (between coordinates 32,182,681-32,185,338) into a p15A minimal vector at a position with flanking loxP and lox511 sequences in the same relative orientations as in the RMCE acceptor site. The 24 PP7 repeats were synthesized (GenScript) along with *Hba-a1* coding sequences between two KflI sites at the 5’ end of the gene and subsequently exchanged by standard restriction enzyme digestion and re-ligation with sequence in the above p15 plasmid to obtain the final p15A-loxP-*Hba-a1*-PP7-lox511 construct to be used as the RMCE donor vector.

### Cell culture

E14-TG2a.IV mouse embryonic stem cells were cultured and differentiated as described previously (Francis et al., 2020, Jeziorska et al., 2017, Nichols et al., 1990, Smith, 1991). Embryoid bodies were harvested and cells released by trypsin-mediated disaggregation. Cell pellets were visually assessed for haemoglobinisation. Cytospins were processed by May-Grünwald-Giemsa (MGG) staining.

### Generation of Hba-a1-PP7 cell line

E14-TG2a.IV mouse embryonic stem (ES) cells were electroporated with the p15A-A22BAC-loxP-pgk-HygTk-lox511 construct to first create the RMCE acceptor site in the place of the *Hba-a1* gene. Electroporated ES cells were selected using hygromycin and resistant clones were analysed with Southern blotting and sequencing of PCR-amplified recombined junctions to identify those arising from correct construct integration by homologous recombination. Next, correctly integrated ES cell clones were co-electroporated with the p15A-loxP-*Hba-a1*-PP7-lox511 plasmid (the RMCE donor vector) and a plasmid expressing Cre recombinase (pCAGGS-Cre-IRESpuro). Cells were selected using ganciclovir to recover those that had undergone RMCE. A ganciclovir resistant clone was validated using Southern blot analysis with probes located at the 5’ and 3’ ends of the locus and with probes corresponding to PP7 repeats and the HygTk cassette. Targeting of a single allele was confirmed by DNA FISH. The presence of PP7 repeats in the locus was also confirmed using MinION Technology sequencing (Oxford Nanopore Technologies). The *Hba-a1*-PP7 ES cell clone was differentiated to EBs, and normal expression and chromatin landscape of the locus was confirmed by RT-qPCR and ATAC-seq.

A pPGK-PCP-GFP-IRES-Hyg plasmid (a kind gift from Jonathan Chubb) designed to constitutively express a PP7 coat protein (PCP)-GFP transgene, was then transfected into unmodified ES cells and into the *Hba-a1*-PP7 clone above. Cells were selected in hygromycin to create stable transgene-expressing cell lines. Those cell lines with a medium and relatively uniform transgene expression level as assessed by GFP fluorescence across the cell population were picked and transgene integration validated by Southern blot analysis.

### Reverse transcription quantitative real-time PCR

RNA was isolated from cells using TRI Reagent (T9424, Merck) before DNase treatment using DNA-free DNA removal kit (AM1906, Thermo Fisher Scientific). Superscript III First-Strand Synthesis SuperMix (11752050, Thermo Fisher Scientific) was used to generate cDNA. qPCR was performed using Fast SYBR Green Master Mix (4385616, Thermo Fisher Scientific) with primers listed in Supplementary Table 1. Data were first normalised to 18S ribosomal RNA at each day of differentiation and then to the maximum value for that gene within the differentiation time series.

### ATAC-seq

ATAC-seq was performed as previously described (Buenrostro et al., 2013). Embryoid bodies were disaggregated at day 7 of differentiation and CD71^+^ Ter119^+^ cells were selected for using magnetic column purification (Miltenyi). 75,000 cells were used per biological replicate. After tagmentation, the DNA was eluted using MinElute columns (28206, Qiagen). PCR indexing was performed using NEBNext High-Fidelity 2x PCR Master Mix (M0541S, NEB) and sequenced using a NextSeq platform (Illumina). After sequencing, read quality was assessed using FASTQC. Data were then aligned to mm9 build of the mouse genome using a custom-built pipeline, where PCR duplicates and ploidy regions were removed, while mitochondrial DNA was excluded during normalization (Telenius et al., 2018).

### DNA FISH

Targeting of PP7 loops to a single allele at the α-globin locus was confirmed using RASER-FISH, a non-heat-denaturing method of DNA FISH, as described previously (Brown et al., 2018). Briefly, cycling cells were grown on coverslips in BrdU/C-containing medium overnight to allow incorporation during DNA replication. Cells were fixed and permeabilised before using exonuclease III (M0206L, NEB) digestion to enable resection of a single DNA strand after treatment with Hoechst 33258 and 254 nm UV light to induce nicks in the BrdU/C-containing DNA strand. Following overnight hybridisation at 37°C and stringency washes to remove mismatched and unbound probes, hybridised probes, where applicable, were detected with appropriate antibodies and nuclei were stained with DAPI before mounting. FISH probes used were ULS550-labelled oligos (FLK 004, Kreatech Biotechnology) against the PP7 repeat sequence (AATTGCCTAGAAAGGAGCAGACGATATGGCGTCGCTCCCT and AGCAGAGCATATGGGCTCGCTGGCTGCAGTATTCCCGGGT) while the 3’ α-globin locus probe was probe ‘pA’ as described in Brown et al. (2018), which was labelled with digoxygenin (DIG) and detected with sheep anti-DIG FITC (11207741910, Roche, RRID: AB_514498) and rabbit anti-sheep FITC antibodies (FI-6000, Vector Laboratories, RRID: AB_2336218). Widefield fluorescence imaging was performed on a DeltaVision Elite system (Applied Precision) equipped with a 100x/1.40 NA UPLSAPO oil immersion objective (Olympus), a CoolSnap HQ2 CCD camera (Photometrics). Filter sets were as follows: DAPI – excitation 390/18, emission 435/40, FITC – excitation 475/28, emission 525/45, TRITC – excitation 542/27, emission 593/45. 12-bit image stacks were acquired with a z-step of 150 nm giving a voxel size of 64.5 nm x 64.5 nm x 150 nm.

### Flow cytometry

Differentiation progression was typically assessed using anti-CD71 PE (113807 Biolegend, RRID: AB_313568) and anti-Ter119 APC (116212 Biolegend, RRID: AB_313712) antibodies at a dilution of 1:10,000 and 1:100 in FACS buffer (1x PBS, 10% FBS) respectively. Hoechst 33258 (1:10,000, H3569, Thermo Fisher Scientific) was used as a Live/Dead marker. Regular flow cytometry experiments were done on an Attune NxT Analyser (Thermo Fisher Scientific), while sorting experiments were done using a FACS Aria Fusion sorter (100 μm nozzle width; BD Biosciences). Day 7 (chosen to ensure a full range of differentiation states) EB cells for sorting were placed in a recovery medium (phenol red-free IMDM base medium, 10% FBS, 1 U/ml erythropoietin) for 1 hour following disaggregation before being stained (anti-CD71 Brilliant Violet 421, 1:5000 dilution – 113813 Biolegend, RRID: AB_10899739; anti-Ter119 Alexa Fluor 647, 1:1000 dilution – 116218 Biolegend RRID: AB_528961) in FACS buffer and sorted based on CD71 and Ter119 signal (Fig. 3B, Extended Data Fig. 5B). A minimum of 250,000 cells were collected for each of 6 sorting gates, with half taken for cytospins and the remainder used for fluorescence microscopy.

### Live imaging

Prior to imaging, embryoid bodies were disaggregated as described above and cells were placed in recovery medium for 4 hours. Cells were then passed through a cell strainer, counted and 500,000 cells allowed to settle on poly-L-lysine coated 35 mm high µ-Dishes (81158, Ibidi) in fresh recovery medium. Cells were then imaged using a 488 nm laser (100 mW) with 100 ms exposure at 50% power every 10 seconds for 10 min for short movies, or every 2.5 min or 5 min for 1 hour for long movies. For measurement of CD71 and Ter119, cells were stained for 5 min before imaging with directly conjugated anti-CD71 Brilliant Violet 421 and anti-Ter119 Alexa Fluor 647 antibodies respectively. Sodium azide was removed from antibodies before use by diluting in 2 ml PBS and re-concentrating using a protein concentrator column (88521, Thermo Fisher Scientific) as this is critical for live imaging experiments using antibodies (Eilken et al., 2009). Multiple stacks were collected in parallel after 1 hour of imaging using 405 nm (50 mW), 488 nm, and 635 nm (30 mW) lasers with 100 ms exposure and at 25%, 50% and 50% power respectively as well as a brightfield DIC image stack. FACS-sorted cells were imaged in the same way but without 1 hour time-lapse imaging. For nuclear labelling experiments, cells were stained with SiR-DNA (SC007, Spirochrome) at 1:1000 dilution for 1 hour. Cells were imaged at 37°C on an inverted Zeiss Cell Observer Spinning Disc system with a CSU-X1M 500 Dual Cam spinning disc unit (Yokogawa), a 1.2x magnification camera C-mount adapter, an Orca-Flash4.0 v2 sCMOS camera (Hamamatsu), and a Plan APO 63x/1.40 NA Oil M27 objective (Carl Zeiss AG) with a final voxel size of 86 nm x 86 nm x 500 nm. Five independent live imaging experiments were performed as described above.

### Image analysis

Fluorescence microscopy images were uploaded to OMERO (Allan et al., 2012) for storage and initial cell identification. Healthy erythrocyte lineage cells were manually identified by morphology: spherical cells lacking obvious blebs, an intact nucleus (mitotic cells were excluded), and absence of intracellular staining with Ter119 antibody (as this was identified as a marker of dead or dying cells). Cell centroids were marked using the ‘point’ ROI tool in OMERO.iviewer and locations and images downloaded from the OMERO server using the Matlab (Mathworks) API.

Manual quantification of spot intensities from cells at day 5, 6 or 7 of embryoid body culture (Extended Data Fig. 4) was done using Imaris (version 9.1.2, Oxford Instruments). Intensity of transcription spots were quantified using the ‘Spots’ tool calculated as the mean of pixels within an ovoid shape of 1 μm x 1 μm x 3 μm in size (x,y,z dimensions respectively), centred on the transcription spot. The nuclear background was measured within a region of size 3 μm x 3 μm x 3 μm within the nucleus at the same z-position but situated away from the transcription spot in the xy plane. Corrected spot intensity was calculated by subtraction of background values from measured spot intensity. If no spot was easily visible (if the gene was inactive) measurements were taken from the coordinates of the last visible spot. Kymographs of SiR-DNA-labelled *Hba-a1-*PP7 nuclei were created in Imaris by first creating a surface for individual channels using simple thresholding. Cells maintaining active transcription spots throughout the imaging period were chosen for ease of visualisation. Single 2D image slices (xy dimensions) from each time point, centred on the transcription spot in the z dimension, were assembled into a 3D image stack with time as the third dimension in order to demonstrate continued nuclear localisation of transcription spots during imaging.

Automated identification and quantification of transcription spot intensities was done in Matlab (version R2014a), largely as described previously (Corrigan et al., 2016), with minor modifications. All other analyses in Matlab were done using version R2020a. In brief, since we did not use a marker such as a fluorescent histone tag to outline the nucleus and given that some MCP-GFP foci were visible in the cytoplasm of some cells, we generated a pseudo-nuclear marker by exploiting the relative depletion of GFP signal in the nucleus compared to cytoplasm (Extended Data Fig. 3B). Cells were analysed individually by using centroid locations to crop the original images, and a simple two-step k-means clustering algorithm was used to segment first the cell boundary, and then the nuclear boundary within this. This estimate of nuclear position was then passed with GFP image stacks to a spot detection algorithm (Corrigan et al., 2016) for automatic detection and measurement of transcription spots. Manual inspection and correction of spot locations was done, where necessary, to ensure high accuracy. This semi-supervised automated analysis showed good agreement with manual quantification of spot intensities in initial experiments (Extended Data Fig. 4). Measurement of CD71 and Ter119 marker levels was done by similarly cropping and segmenting a cell mask from the GFP channel, before taking the mean value in each channel of all pixels within the mask, minus an estimated background (iteratively smoothed from pixels outside the cell mask until no change is observed, as described previously, Corrigan et al., 2016) for each cell. Cell size was approximated by taking the area of the cell mask in the central z-slice (centroid of the mask in the z-dimension) for each cell. Given the highly spherical nature of early erythroid cells, this proved to be a sufficiently good estimate for our studies.

MGG cytospins were scored manually according to changes in size, colouration and texture of nuclei and cytoplasm of cells using the ‘Cell Counter’ plugin in Fiji (Schindelin et al., 2012) after image names were randomized. Erythroblast counts for each image were then unblinded and collated according to FACS-sorted populations (F1-F6).

### Data analysis

Short movies (10 min; Extended Data Fig. 6) were used to establish an optimum frame rate for longer-term imaging of transcription dynamics (1 hour). Transcription spot intensities were extracted from images as described above. The data were then subsampled at progressively increasing frame intervals for each cell to simulate use of different experimental imaging sampling intervals. We wanted to establish the optimum sampling interval by which to capture all transcriptional bursting events while also minimising light input and associated photobleaching. To do this, we estimated the amount of information loss which occurs with increasing frame rate by calculating the root mean-squared error (RMSE) when comparing the raw data (sampled every 10 s, *y_raw_*) to subsampled data (*y_sub_*) for each cell:

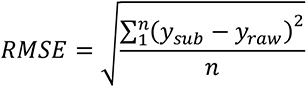

A lower RMSE indicates reduced error (information loss) of subsampled compared to raw data. A sampling interval of 150 s was found to be a good compromise to minimise both light input and information loss.

Differentiation progression for cells imaged by microscopy was estimated using the measured intensities of CD71 and Ter119 staining (Extended Data Fig. 7). Cells undergoing embryoid body differentiation exhibit known changes in the levels of these two markers during differentiation progression, from CD71^-^/Ter119^-^, to CD71^+^/Ter119, to CD71^+^/Ter119^+^ (Francis et al., 2020; Extended Data Fig. 5A). We used this knowledge to measure the progression of each cell through this differentiation space. To do this, we empirically defined a series of curves that followed this pattern of changes in CD71 and Ter119 markers. Several curves were used in order to capture the full range of potential trajectories through this CD71/Ter119 axis, given that a range of CD71 intensities were observed in Ter119^low^ cells. These curves were then discretised into 10,000 equally spaced points using the ‘interparc’ function in Matlab. We measured the distance between each cell and every point on these curves, found the point closest to each cell, and calculated the proportional distance of that point along its curve as a fraction of 1. This value was taken as a proxy for the extent of differentiation progression for each cell.

Having defined a series of differentiation stages (DS1-DS6) from time-lapse imaging long movies we wanted to estimate which erythroblast stages were most likely to represent these stages. To do this, we used microscopy measurements of CD71 and Ter119 intensity to estimate differentiation progression (as above) of day 7 EB FACS-sorted cell populations (F1-F6) (Fig. 3B, Extended Data Fig. 7F). Unsorted day 7 cells (Extended Data Fig. 5A) were also included to ensure enough cells to enable comparison to the long movie time-lapse dataset. To allow comparison between the time-lapse dataset and the FACS-sorted dataset we used regular fluctuations in the size of cells along the differentiation axes (from 0 to 1) to align the datasets in differentiation. Following this alignment, the coordinates of cells in differentiation should be equivalent for DS1-DS6 and F1-F6. Given that we assessed the identity of erythroblast stages in F1-F6 (Fig. 3C, D), and now knowing where these cells were on the differentiation axis, we could approximate the cell identity in time-lapse imaging differentiation stages. Firstly, the distribution of FACS-sorted cells on the differentiation axes were plotted separately for each sorted population F1-F6 (Extended Data Fig. 7Gi). Secondly, the known proportions of erythroblast stages for each of these populations as assessed by morphology analysis (Fig. 3C, Extended Data Fig. 7Gii) were overlaid onto these histogram distributions from least to most differentiated (proerythroblast, basophilic erythroblast, polychromatic erythroblast, orthochromatic erythroblast; Extended Data Fig. 7Giii). Finally, using the aligned coordinates of the time-lapse imaging differentiation stages (e.g. DS1 at 0 to ∼ 0.1) on this differentiation axis we assigned erythroblast cell identity to each of these groups by visual assessment of the erythroblast stages across F1-F6 at these coordinates (Extended Data Fig. 7Giii).

Confidence intervals for the median of mean spot intensities for cells at each differentiation stage (DS1-DS6) were calculated using bootstrapping (Extended Data Fig. 8B). For each differentiation stage, the distribution of mean spot intensity values was randomly sampled with replacement (n = 31, to match the lowest number of cells in any differentiation stage, DS1) 10,000 times and median values determined. Limits of 95% confidence intervals were calculated as the 2.5^th^ and 97.5^th^ percentiles of this bootstrapped distribution.

The threshold above which α-globin was considered to be active or ‘ON’ was defined empirically by visual inspection of images to be 350 arbitrary intensity units. However, at each analysis stage, care was taken to test a number of thresholds around this value to ensure that conclusions are consistent regardless of which ON/OFF threshold was chosen. The ON fraction was calculated for each cell as the fraction of time the gene spends in the ON state as a proportion of the total imaging period. To more precisely estimate time spent above the ON/OFF threshold, we used linear interpolation of individual transcription traces. Duration of individual bursts was measured similarly but only ‘complete’ bursting events were included, where both the start and end of a burst was below the ON/OFF threshold. Transcriptional output of an individual cell was calculated as the area above the ON/OFF threshold and below the fluctuating (interpolated) transcriptional activity trace summed across the imaging period. This is effectively the summed burst size for all bursting events (including those ‘incomplete’ bursts). Burst size for individual bursts was measured similarly but again only included ‘complete’ bursting events. Calculating the number of complete bursts identified across a large range of ON/OFF thresholds demonstrated a peak centred around our empirically defined intensity threshold of 350, further supporting the use of this threshold estimate (Extended Data Fig. 10B).

In order to categorise cells according to differing transcriptional behaviours, we first distinguished cells according to the length of time spent in the active state (ON duration). We did this based on the relationship between the ON duration and transcriptional output which we used robust LOWESS local regression (‘rlowess’ in Matlab ‘smooth’ function) to model. A clear inflection point suggested differences in burst amplitude at high ON duration (see Supplementary Note) and we therefore categorised cells as ‘High ON’ or ‘Low ON’ according to whether their ON duration was above or below this inflection point, respectively (Fig. 5A). We then categorised cells according to the amplitude of transcriptional bursts displayed over the imaging period as we noticed many cells consistently showed a low or ‘basal’ level of burst amplitude while some exhibited much higher burst amplitude. We again used the relationship between ON duration and transcriptional output to define a threshold above which cells would be classed as exhibiting ‘high amplitude bursting’ on average across the imaging period (Fig. 5C). Firstly, we estimated the relationship between ON duration and transcriptional output for a basal amplitude bursting regime by extrapolating from the local regression described above for Low ON cells only, as the majority of these cells exhibited basal amplitude bursting. Fitting a quadratic curve to the Low ON regression data points allowed us to estimate the transcriptional output for basal amplitude bursting in High ON cells. To categorise cells according to whether they exhibit ‘basal’ or ‘high’ amplitude bursting, we then wanted to measure how far each cell deviates from basal amplitude bursting, in terms of its transcriptional output. If the transcriptional output of a particular cell is very close to what would be expected from basal amplitude bursting with a certain ON duration then this deviation would be small. In contrast, for cells exhibiting high amplitude bursting, and therefore with a high transcriptional output for a particular ON duration, this deviation would be much higher. To more easily define a single threshold value with which to identify high amplitude bursting cells, we first calculated the residual between the transcriptional output of each cell and the basal amplitude estimate of this value for the same ON duration. We then normalised these residuals by the ON duration, to account for the increased time for higher ON cells to deviate from a basal amplitude bursting regime. Finally, we used the distribution of these normalised ‘amplitude residuals’ to set the high amplitude bursting threshold. In Low ON cells, as expected we found a close-to normal distribution of amplitude residuals around a value of 0 (meaning the deviation of most of these cells from ‘basal amplitude bursting’ was small), with a skewed tail of high amplitude bursting cells. Setting the threshold as one standard deviation away from the median enabled convenient separation of basal and high amplitude cells. Cells were then classed into three groups defined by these thresholds and indicative of differing transcriptional behaviours: 1. Basal amplitude (both Low and High ON), 2. High ON, High amplitude, 3. Low ON, High amplitude.

## Acknowledgements

We are very grateful to Jonathan Chubb for helpful comments on the manuscript; Damien Downes for assistance with submission of data to the Gene Expression Omnibus; Dominic Waithe for initial suggestions for image analysis; Craig Waugh and the Flow Cytometry Facility at the Weatherall Institute for Molecular Medicine (WIMM) for help with FACS sorting experiments; the Computational Biology Research Group and Wolfson Imaging Centre at the WIMM for bioinformatic support and use of microscopy facilities and associated services. This work was supported by Medical Research Council grants to D.R.H. (MC_UU_00016/4, MR/T014067/1).

## Data availability

Data presented in this work are available from the authors upon request. ATAC-seq raw data files are available from the Gene Expression Omnibus (GEO) at accession number GSE189474.

## Code availability

Image analysis was primarily done using published Matlab code (Corrigan et al., 2016, see http://www.ucl.ac.uk/lmcb/sites/default/files/Corrigan2016MatlabFiles.zip). All other code for this work is available from the authors upon request.

---

**Extended Data Figure 1:**
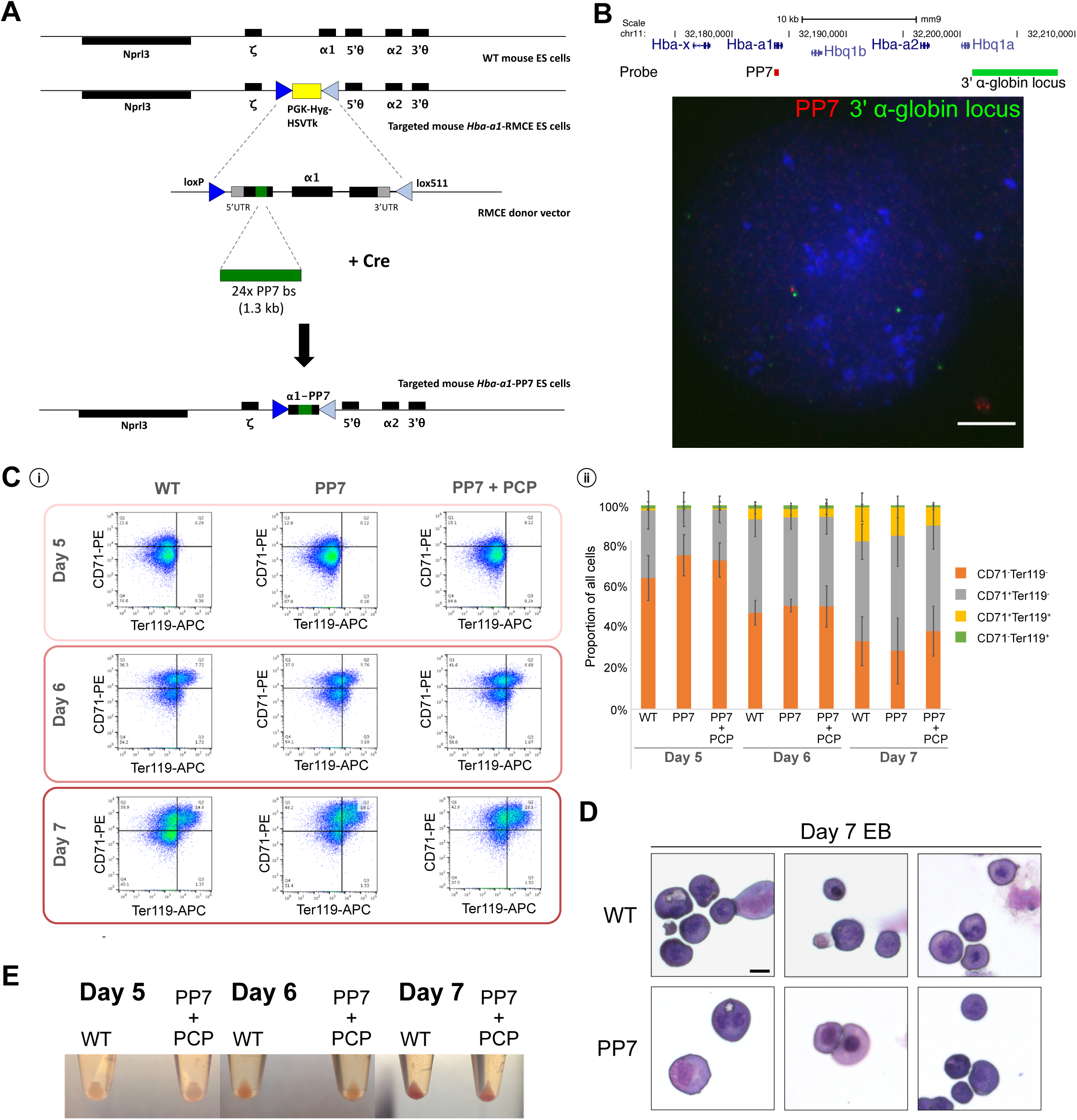
Generation and characterisation of *Hba-a1*-PP7 cell line. (A) A array of 24 bacteriophage PP7 repeats was integrated into the first exon of *Hba-a1* in mouse E14 ES cells using an RMCE system. (B) Representative example of DNA FISH indicating colocalization of the PP7 repeats (red) with a probe sited at the 3’ end of α-globin locus (green) in targeted mES cells. Scale = 5 μm. (C) Representative flow cytometry plots of CD71 vs Ter119 staining during EB differentiation (i) and quantification of proportion of cell population within each gate (ii). Changes in CD71 and Ter119 markers during differentiation are equivalent in WT and *Hba-*a1-PP7 (+PCP) cells. Error bars = standard deviation. (D) Examples of May-Grunwald-Giemsa-stained EB cells (WT and *Hba-a1*-PP7) at day 7 of differentiation. A range of erythroblast stages are observed in both cell lines. Scale = 10 μm. (E) Disaggregated EB cell pellets from WT E14 mES and *Hba-a1*-PP7 + PCP-GFP cells showing equivalent haemoglobinisation during differentiation.

**Extended Data Figure 2:**
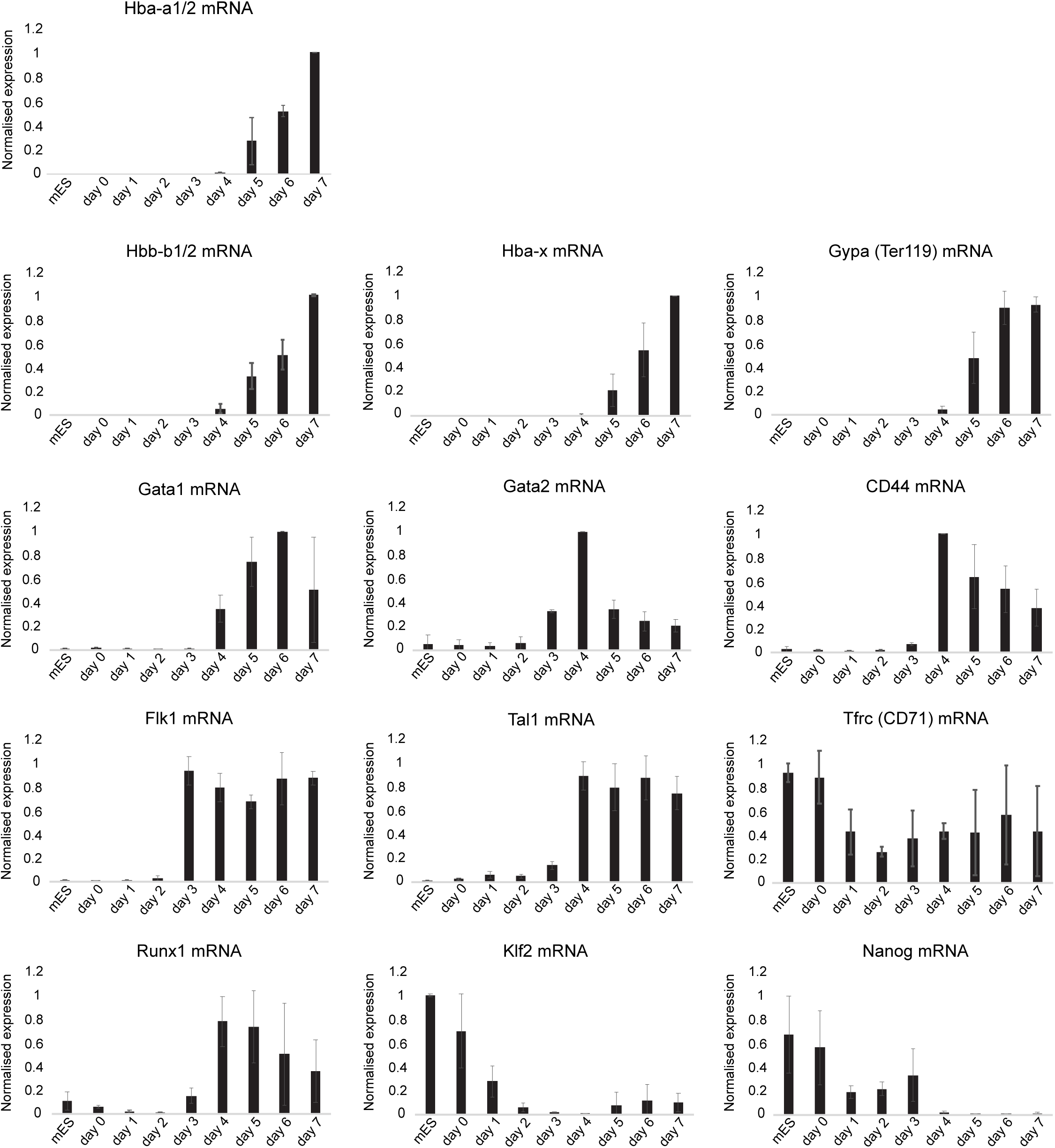
RT-qPCR time course of EB differentiation from WT mES cells. A panel of stem cell and erythroid markers were profiled at each day of differentiation from mES cells to EBs. Data are normalised to Rn18s and then within differentiation time course for each gene. Mean of three independent experiments shown. Error bars = standard deviation.

**Extended Data Figure 3:**
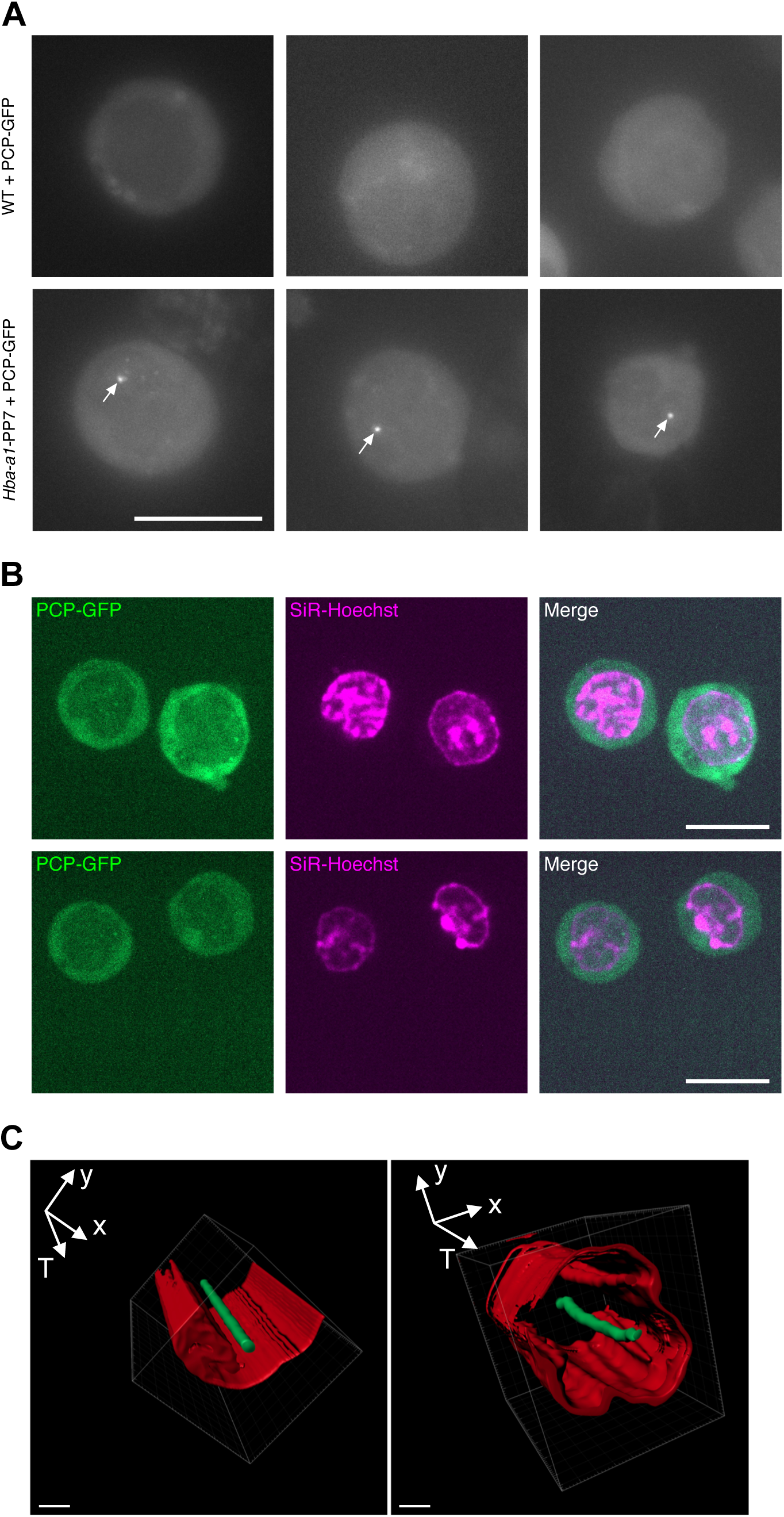
Nuclear-localised foci in *Hba-a1*-PP7 cells represent sites of active transcription. (A) Examples of cells expressing PCP-GFP in WT or *Hba-a1*-PP7 clones. Bright nuclear localised foci (arrows) are only detected in cells containing PP7 loops in the *Hba-a1* gene, indicating ongoing transcriptional activity. Scale = 10 μm. (B) Co-staining of *Hba-a1*-PP7 cells with SiR-Hoechst shows depleted PCP-GFP intensity within the nucleus, enabling identification of nuclear transcription spots. Scale = 10 μm. (C) Imaris software reconstruction of representative cells over time, T, showing that bright nuclear foci (green, PCP-GFP) stay within the nucleus (red, SiR-Hoechst) for the duration of the experiment. Scale = 2 μm.

**Extended Data Figure 4:**
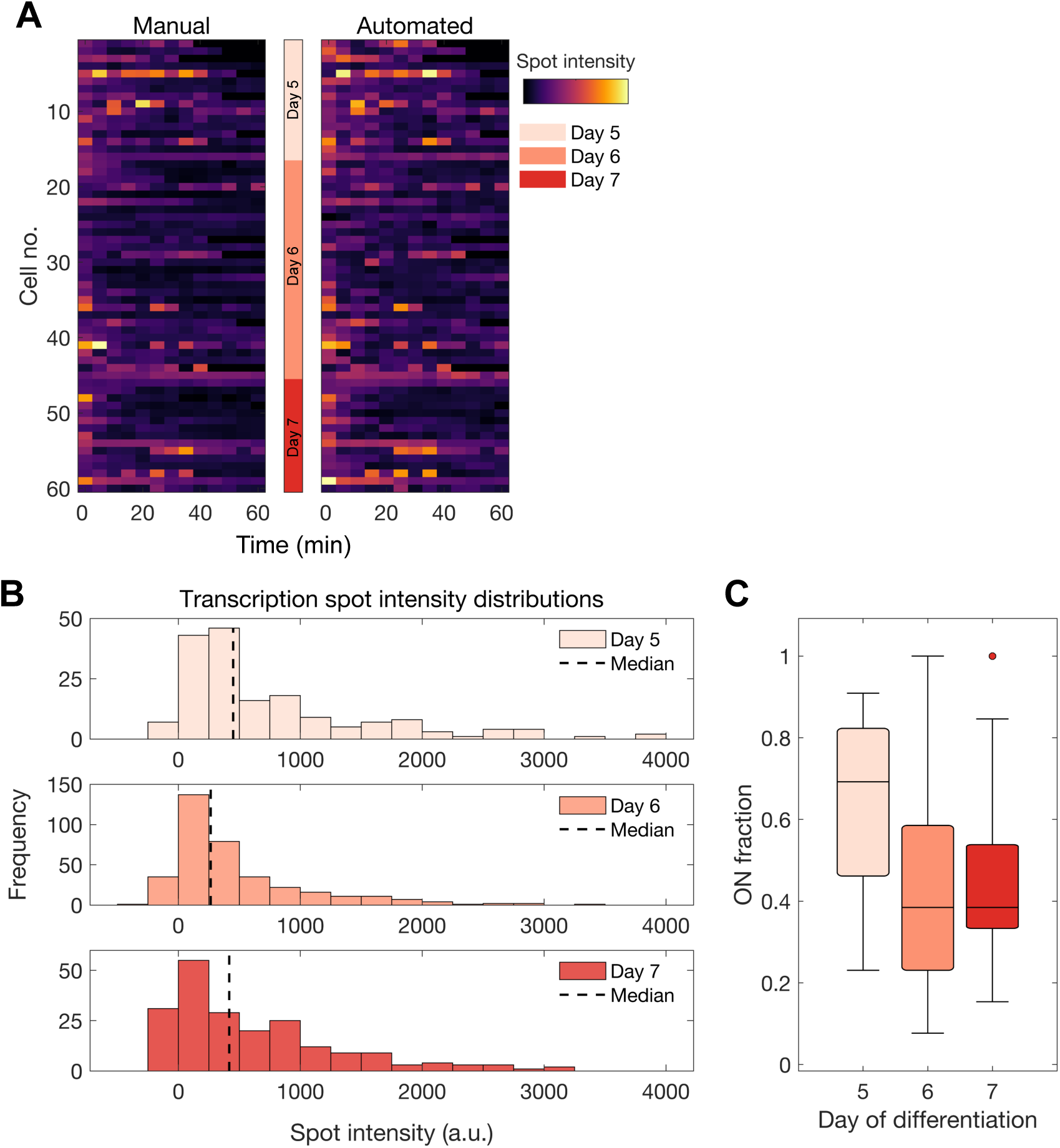
α-globin transcription dynamics at day 5, 6 and 7 of EB differentiation. (A) Spot intensity traces of 60 cells from day 5, 6 and 7 of EB differentiation imaged for 1 h at 5 min frame rate. Each row represents a single cell. Cells were analysed both manually and using a semi-automated approach and the results are highly similar. (B-C) Distributions of transcription spot intensities (B) and fraction of time spent active (C) for cells cultured for 5, 6, or 7 days. Boxes within boxplots show median and interquartile range, whiskers show 9^th^ and 91^st^ percentile of distribution. a.u. = arbitrary units.

**Extended Data Figure 5:**
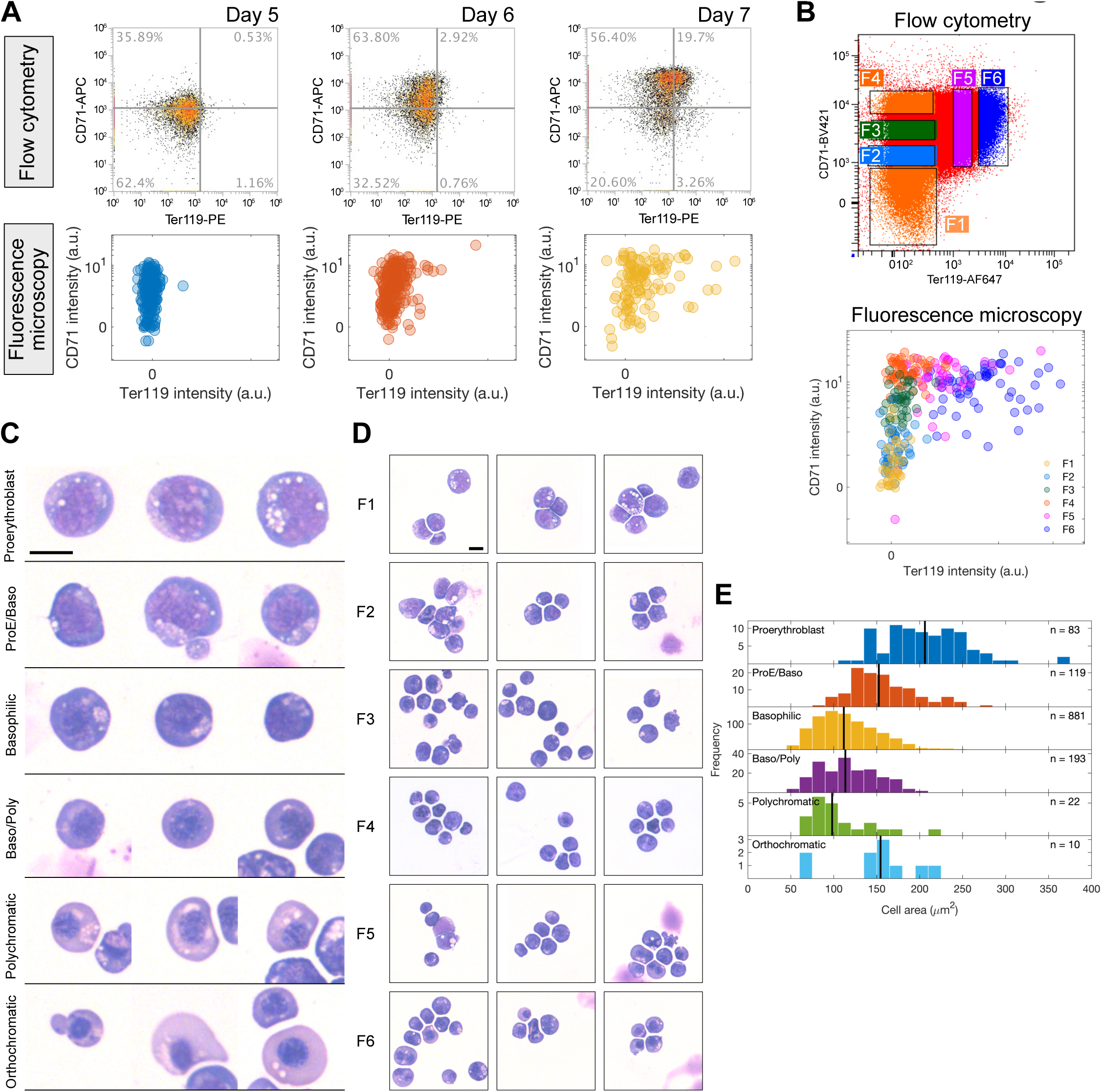
Validation of on-microscope staging of EB-derived erythroid cells. (A) Comparison of CD71/ Ter119 staining patterns and analysis of cells when measured by either flow cytometry or microscopy. Cells were derived from the same culture and sampled throughout a differentiation time course. (B) Day 7 EB cells were sorted into 6 populations (F1-F6) and subsequently imaged, demonstrating equivalent segregation of cells using the two modalities. (C) FACS-sorted cells from (B) were also stained with May-Grunwald-Giemsa (MGG) stain and scored according to morphology into 6 erythroblast stages, with representative example cells shown for each class. (D) Representative fields of view of MGG staining for each FACS-sorted population (F1-F6). (E) Cell area distributions for each erythroblast class. Solid line shows median cell area. n = number of cells scored. Decreasing size of progressive stages of differentiation is a well-known feature of erythropoiesis, validating our approach.

**Extended Data Figure 6:**
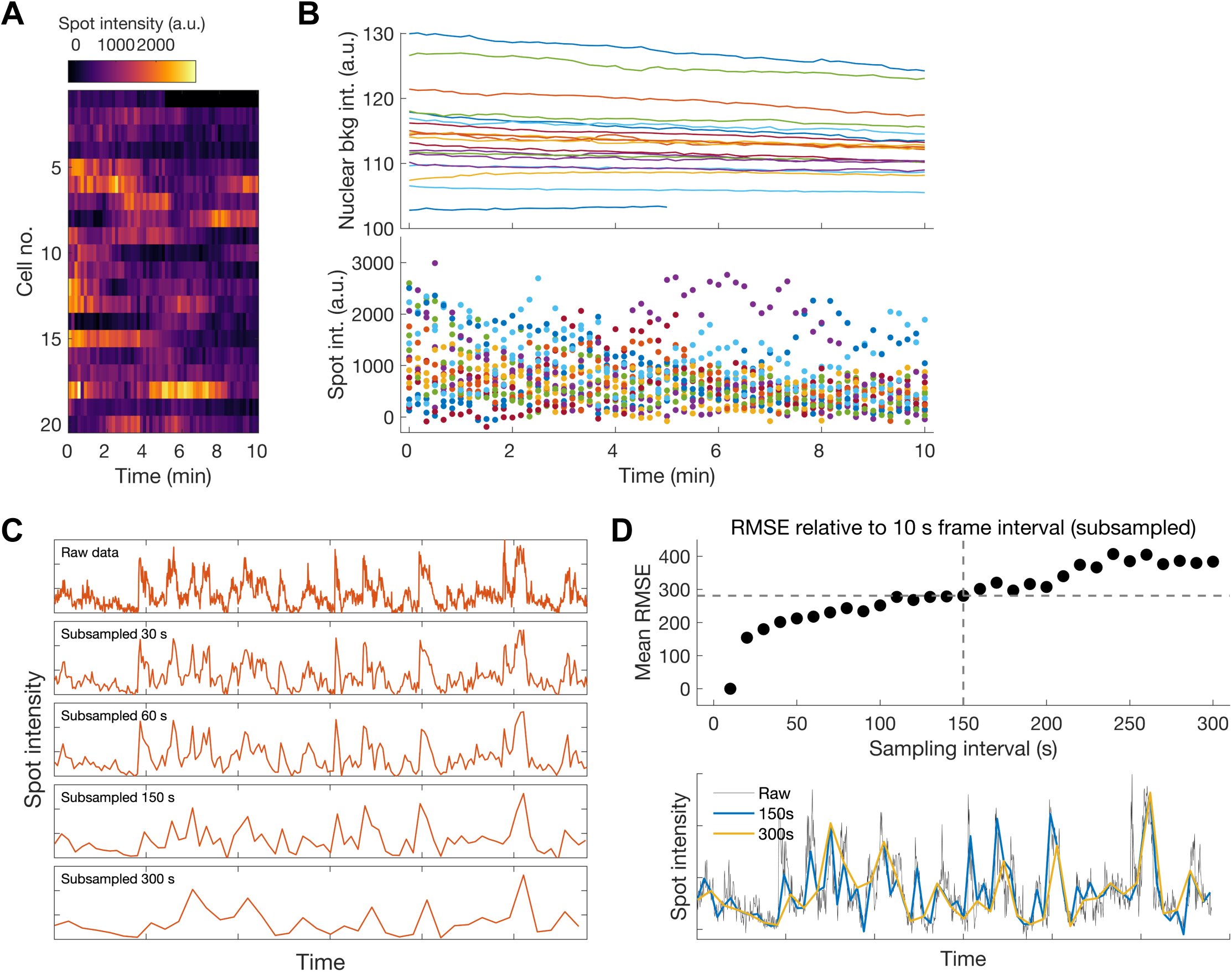
Minimising photobleaching during live imaging experiments. (A) Heatmap of transcription spot intensity time series of cells imaged at high frame rate (every 10 s) for 10 min. Each row is a single cell. (B) Average nuclear background (top), and individual spot intensities (bottom) measured across image capture sessions. Different colours identify individual cells. Downward slope of overall intensity measurements during the movie indicates photobleaching and is most clearly visible after about 30 frames (5 min). a.u. = arbitrary units. (C) Raw data (top panel) from (A) was concatenated and computationally subsampled at increasing frame intervals to enable estimate of required frame rate for full capture of transcription dynamics. (D) Top panel: root mean squared error (RMSE) estimates of information lost from raw to subsampled data at increasing sampling intervals. A sampling interval of 150 s (2.5 min) represents a compromise to minimise both photobleaching and loss of dynamic transcription information. Bottom panel: visualisation of 2.5 min as an intermediate value enabling minimisation of number of frames to reduce photobleaching, while enabling capture of majority of peaks in the data, compared to 5 min sampling interval.

**Extended Data Figure 7:**
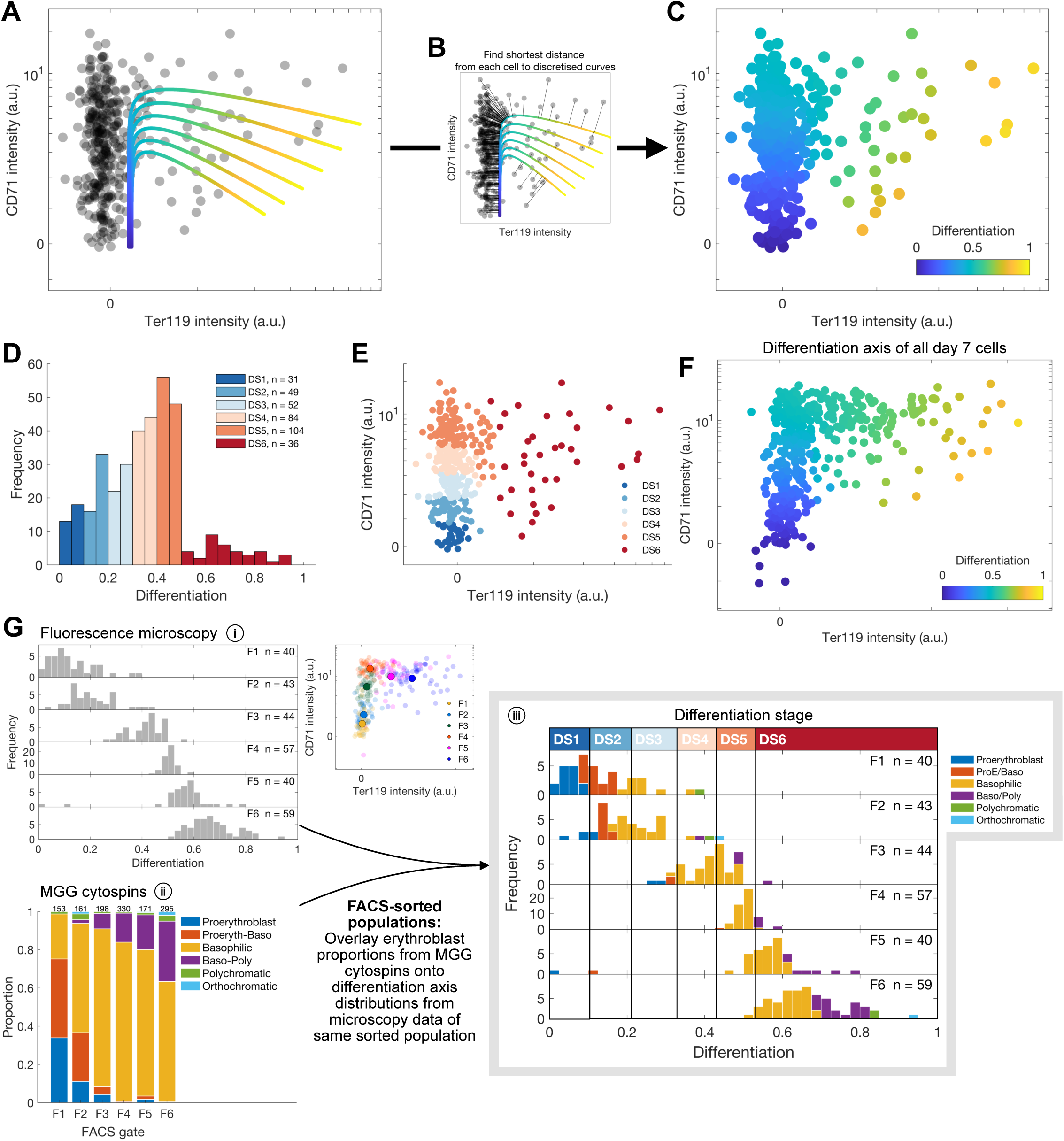
Staging cells in differentiation by changes in cell surface marker intensities. (A) Multiple empirically defined curves following known changes of cell surface marker levels in differentiation (CD71^-^/Ter119^-^, CD71^+^/Ter119^-^, CD71^+^/Ter119^+^, see Ext. Data Fig. 1C, 5A) were overlayed onto the CD71/Ter119 axis. (B) Each curve was discretised into individual points, and each cell was mapped to the nearest point along one of the curves. (C) The fractional distance (scaled from 0-1) of each cell along the curves was taken as a measure of differentiation progression for that cell. Blue cells are early in differentiation, orange/yellow cells are later in differentiation. (D) Histogram of position of cells along the pseudo-differentiation axis. Cells were partitioned into differentiation stages (DS1-6) according to arbitrary lengths of this differentiation axis (0.1 for DS1-DS5 and 0.5 for DS6 due to lower number of cells). (E) Cells in differentiation stages (DS1-DS6) from one-dimensional representation in D mapped back onto two-dimensional CD71/Ter119 axis, as in A. (F) As for Ext. Data Fig. 7C, the position of FACS-sorted cell populations (F1-F6, Fig. 3B) along the differentiation axis was determined (blue = early differentiation, orange = late differentiation) using levels of CD71 and Ter119 intensity (as measured by fluorescence microscopy, Fig. 3B). (G) (i) For each FACS-sorted population (F1-F6), the distribution of cells along the differentiation axis (from 0-1) is shown as a histogram. The proportion of cells from the same sorted populations classified into erythroblast stages by MGG staining (ii) (as in Fig. 3D) were then overlaid onto these distributions from least to most mature (iii). The position of cells in differentiation from time-lapse imaging experiments (DS1-DS6) and FACS-sorted populations (F1-F6) could then be compared. This allowed inference of the erythroblast identity of cells in DS1-DS6. For example, DS1, at 0-0.1 on the differentiation axis, is in line with proerythroblasts (blue) across F1-F6 populations. Similarly, DS6 is a mix of late basophilic (yellow), polychromatic (purple/green) and orthochromatic (cyan) erythroblasts (∼0.5-1 on the differentiation axis).

**Extended Data Figure 8:**
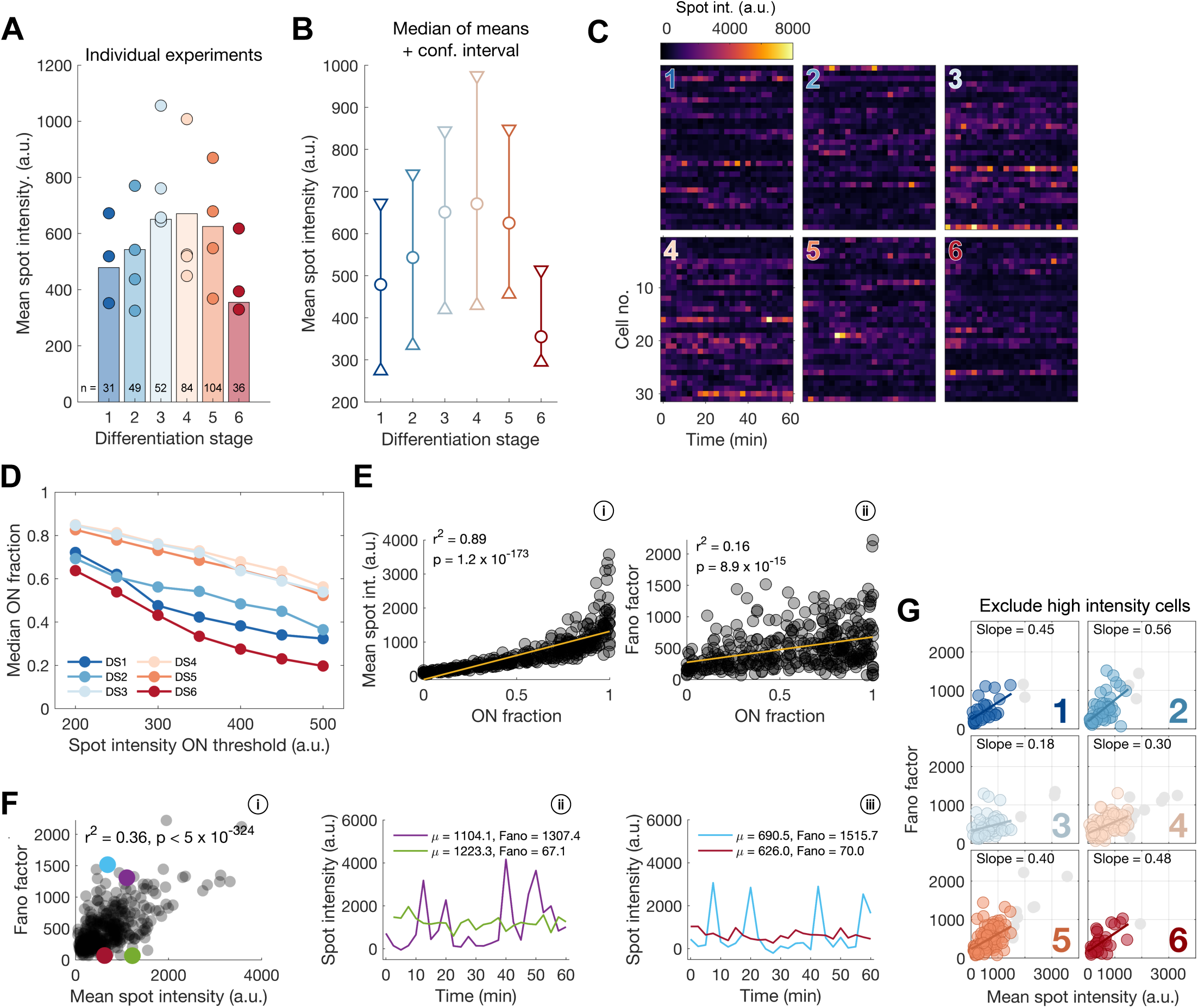
Changes in α-globin transcription dynamics during erythroid differentiation. (A) Median values of mean spot intensity distributions for individual experiments (circles, where n > 5 for that differentiation stage) and all cells (bars). a.u. = arbitrary units. (B) 95% confidence intervals for the median value of the distribution of mean spot intensities for each cell. calculated by bootstrapping (random sampling with replacement) 10,000 times. Observed sample medians shown as circles. (C) Heatmaps of raw spot intensity data for example individual cells across differentiation stages. (D) Median ON fraction for a range of ON/OFF thresholds for each differentiation stage. DS3-DS5 are consistently highest regardless of threshold used. (E) Spearman’s rank correlation of ON fraction with mean spot intensity (i) or Fano factor (ii) for all cells. High correlation between ON fraction and mean spot intensity suggests α-globin transcriptions levels are strongly dependent on time spent transcribing. (F) Relationship between mean spot intensity and Fano factor for all cells (Spearman’s rank correlation) (i), and illustrative examples of individual cell spot intensity traces (ii-iii). (G) Relationship between mean spot intensity and Fano factor across differentiation stages (DS1-DS6, as in Fig. 4E) with highly-transcribing cells removed (greyed out). The trend in the relationship across differentiation is maintained.

**Extended Data Figure 9:**
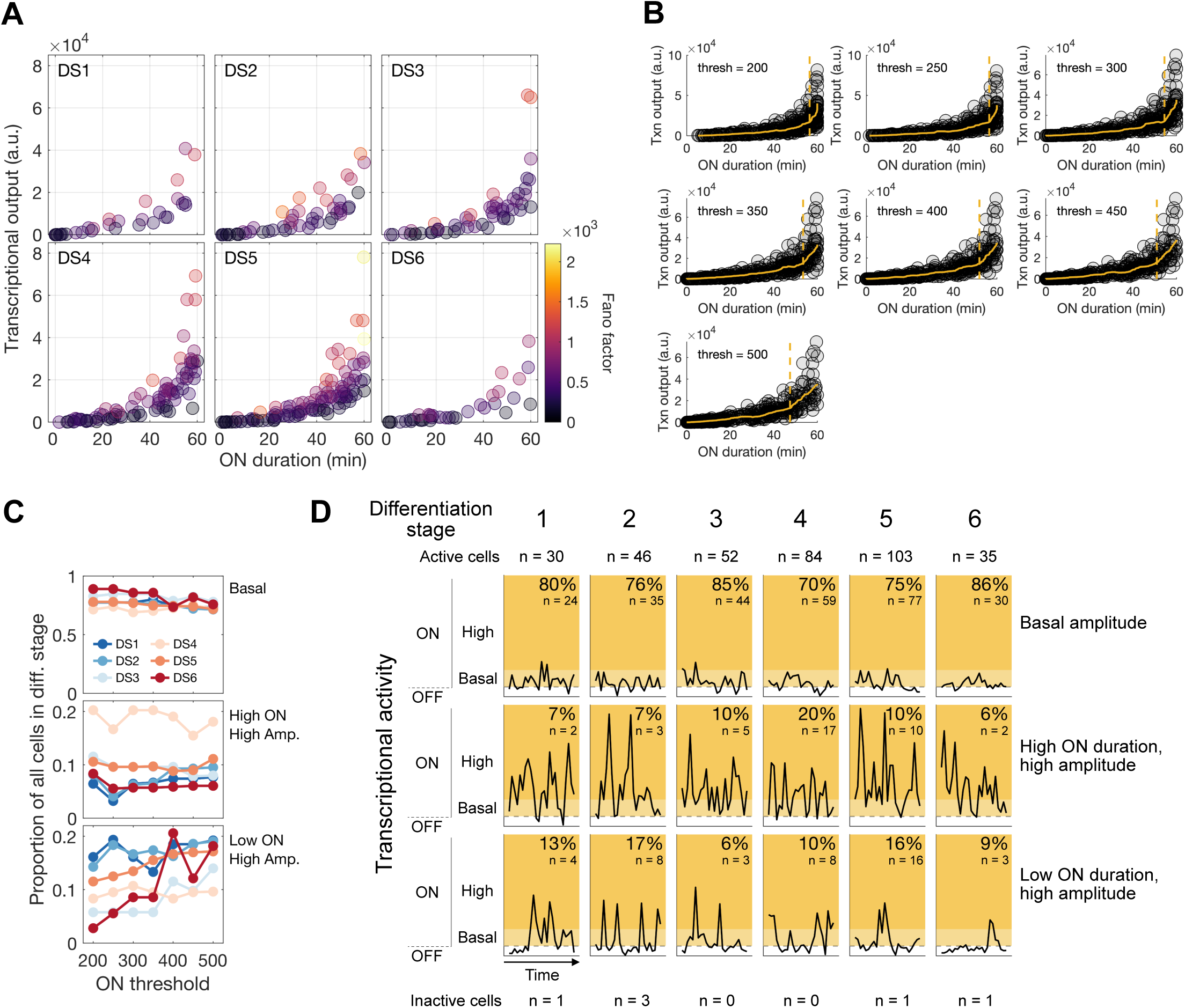
Different α-globin transcriptional behaviours during differentiation. (A) Relationship between total duration of active transcription for each cell (ON duration) and transcriptional output across differentiation stages. Colours indicate transcriptional variability defined by the Fano factor. a.u. = arbitrary units. (B) Identification of Low/High ON duration inflection point to facilitate calculation of different behaviour proportions with varying ON/OFF thresholds. (C) The proportion of different behaviours – basal amplitude transcription, high ON high amplitude transcription, low ON high amplitude transcription – throughout differentiation with varying ON/OFF threshold. ON/OFF threshold does not influence relative proportion of these behaviours at progressive differentiation stages. (D) Representative examples along with summary statistics of the number and proportions of cells with different transcriptional behaviours throughout erythropoiesis. Inactive cells are those which do not display an active transcription spot within the imaging period.

**Extended Data Figure 10:**
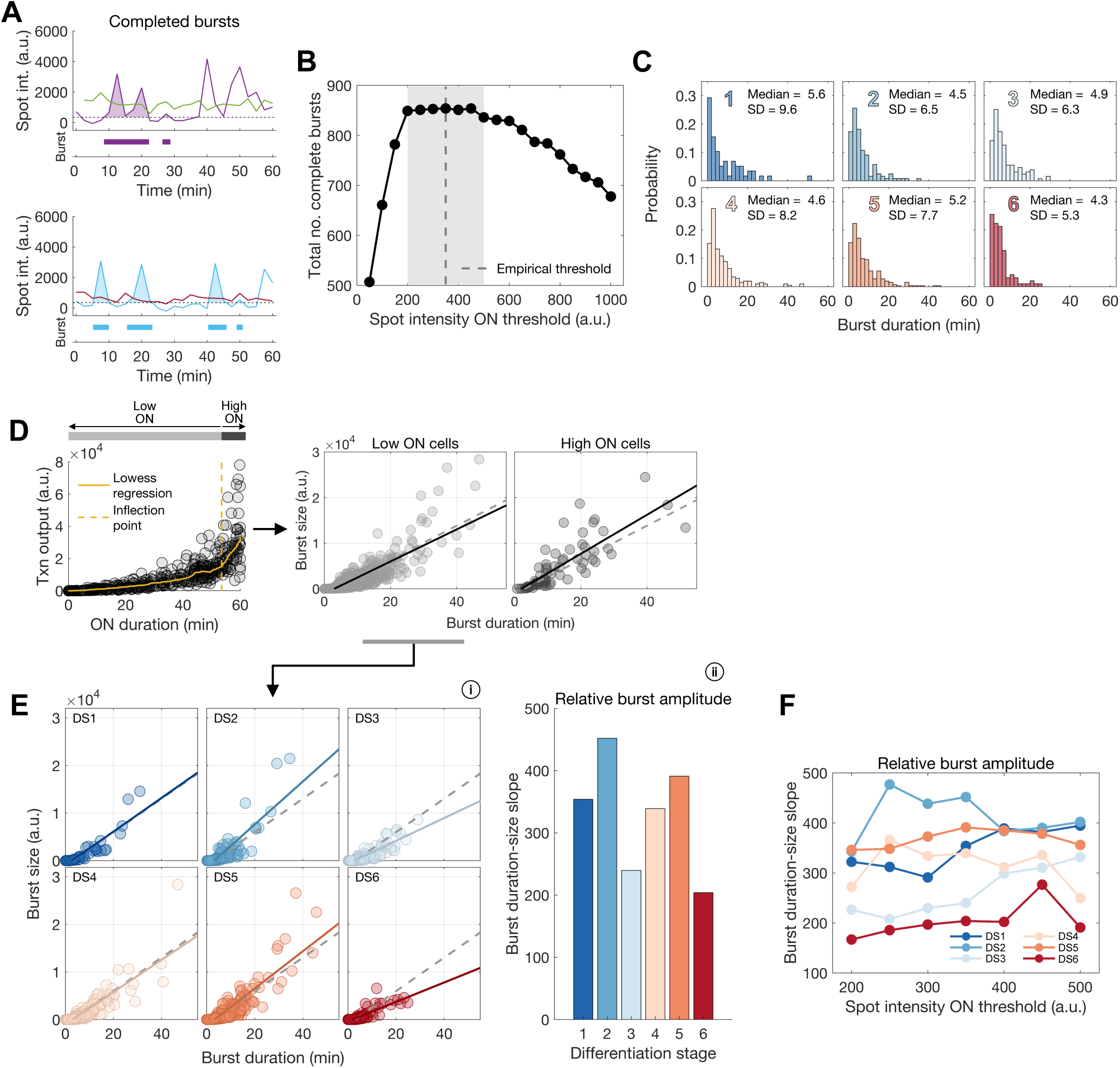
Burst amplitude changes during differentiation (A) Examples of completed bursts within spot intensity traces. Only bursts which begin and end below the ON/OFF threshold are counted. Int = intensity, a.u. = arbitrary units. (B) Total number of completed bursts identified from all cells with varying ON/OFF thresholds. (C) Distributions of individual burst duration for differentiation stages 1-6. (D) Comparison of burst duration-size relationship for low ON and high ON cells derived from Fig. 4A. Dashed line represents linear regression for all cells as reference, solid line for either low or high ON cells. High ON cells have a higher burst size for a given burst duration, and therefore a higher burst amplitude. Txn = transcriptional. (E) (i) Burst duration-size relationship for low ON cells split by differentiation stage, enabling comparison of relative burst amplitude in these cells. Solid lines are linear regression fit of cells in differentiation stages, dashed lines represent fit for all cells as a reference point. (ii) Slope of fits in (i). This represents a relative measure of burst amplitude. (F) Slope of burst duration-size relationship as in (E) with varying ON/OFF thresholds. The trends in terms of relative burst amplitude are maintained across differentiation regardless of ON/OFF threshold.

## References

1. Alexander, J.M., Guan, J., Li, B., Maliskova, L., Song, M., Shen, Y., Huang, B., Lomvardas, S., and Weiner, O.D. (2019). Live-cell imaging reveals enhancer-dependent Sox2 transcription in the absence of enhancer proximity. eLife 8.

2. Allan, C., Burel, J.-M., Moore, J., Blackburn, C., Linkert, M., Loynton, S., MacDonald, D., Moore, W.J., Neves, C., Patterson, A., et al. (2012). OMERO: flexible, model-driven data management for experimental biology. Nature Methods 9, 245–253.

3. Bartman, C.R., Hsu, S.C., Hsiung, C.C.-S., Raj, A., and Blobel, G.A. (2016). Enhancer Regulation of Transcriptional Bursting Parameters Revealed by Forced Chromatin Looping. Molecular Cell 62, 237–247.

4. Boija, A., Klein, I.A., Sabari, B.R., Dall’Agnese, A., Coffey, E.L., Zamudio, A.V., Li, C.H., Shrinivas, K., Manteiga, J.C., Hannett, N.M., et al. (2018) Cell, 175 (7) 1842–1855.e16.

5. Bothma, J.P., Garcia, H.G., Esposito, E., Schlissel, G., Gregor, T., and Levine, M. (2014). Dynamic regulation of eve stripe 2 expression reveals transcriptional bursts in living Drosophila embryos. Proceedings of the National Academy of Sciences 111, 10598–10603.

6. Brouwer, I. and Lenstra, T.L. (2019). Visualizing transcription: key to understanding gene expression dynamics. Current Opinion in Chemical Biology 51, 122–129

7. Brown, J.M., Leach, J., Reittie, J.E., Atzberger, A., Lee-Prudhoe, J., Wood, W.G., Higgs, D.R., Iborra, F.J., and Buckle, V.J. (2006). Coregulated human globin genes are frequently in spatial proximity when active. Journal of Cell Biology 172, 177–187.

8. Brown, J.M., Roberts, N.A., Graham, B., Waithe, D., Lagerholm, C., Telenius, J.M., De Ornellas, S., Oudelaar, A.M., Scott, C., Szczerbal, I., et al. (2018). A tissue-specific self-interacting chromatin domain forms independently of enhancer-promoter interactions. Nature Communications 9, 3849.

9. Buenrostro, J.D., Giresi, P.G., Zaba, L.C., Chang, H.Y. and Greenleaf, W.J. (2013) Transposition of native chromatin for fast and sensitive epigenomic profiling of open chromatin, DNA-binding proteins and nucleosome position. Nature Methods 10 1213–1218.

10. Chao, R., Gong, X., Wang, L., Wang, P., and Wang, Y. (2015). CD71high population represents primitive erythroblasts derived from mouse embryonic stem cells. Stem Cell Research 14, 30–38.

11. Chen, H., Levo, M., Barinov, L., Fujioka, M., Jaynes, J.B., and Gregor, T. (2018). Dynamic interplay between enhancer–promoter topology and gene activity. Nature Genetics 50, 1296–1303.

12. Chubb, J.R., Trcek, T., Shenoy, S.M., and Singer, R.H. (2006). Transcriptional Pulsing of a Developmental Gene. Current Biology 16, 1018–1025.

13. Corrigan, A.M., Tunnacliffe, E., Cannon, D., and Chubb, J.R. (2016). A continuum model of transcriptional bursting. eLife 5, 631.

14. Dar, R.D., Razooky, B.S., Singh, A., Trimeloni, T.V., McCollum, J.M., Cox, C.D., Simpson, M.L., and Weinberger, L.S. (2012). Transcriptional burst frequency and burst size are equally modulated across the human genome. Proceedings of the National Academy of Sciences 109, 17454–17459.

15. Donovan, B.T., Huynh, A., Ball, D.A., Patel, H.P., Poirier, M.G., Larson, D.R., Ferguson, M.L., and Lenstra, T.L. (2019). Live-cell imaging reveals the interplay between transcription factors, nucleosomes, and bursting. The EMBO Journal e100809.

16. Eilken, H.M., Nishikawa, S.-I., and Schroeder, T. (2009). Continuous single-cell imaging of blood generation from haemogenic endothelium. Nature 457, 896–900.

17. Francis, H.S., Harrold, C.L., Beagrie, R.A., King, A.J., Gosden, M.E., Blayney, J.W., Jeziorska, D.M., Babbs, C., Higgs, D.R., and Kassouf, M.T. (2020) Scalable in vitro production of defined mouse erythroblasts. bioRxiv doi: 10.1101/2020.11.10.376749

18. Fraser, S.T., Isern, J., and Baron, M.H. (2006). Maturation and enucleation of primitive erythroblasts during mouse embryogenesis is accompanied by changes in cell-surface antigen expression. Blood 109, 343–352.

19. Fritzsch, C., Baumgärtner, S., Kuban, M., Steinshorn, D., Reid, G., and Legewie, S. (2018). Estrogen-dependent control and cell-to-cell variability of transcriptional bursting. Molecular Systems Biology 14 (2) e7678.

20. Fukaya, T., Lim, B., and Levine, M. (2016). Enhancer Control of Transcriptional Bursting. Cell 166, 358–368.

21. Furlong, E.E.M., and Levine, M. (2018). Developmental enhancers and chromosome topology. Science 361, 1341–1345.

22. Garcia, D.A., Johnson, T.A., Presman, D.M., Fettweis, G., Wagh, K., Rinaldi, L., Stavreva, D.A., Paakinaho, V., Jensen, R.A.M., et al. (2021) An intrinsically disordered region-mediated confinement state contributes to the dynamics and function of transcription factors. Molecular Cell 81 (7), 1484–1498.e6

23. Golding, I., Paulsson, J., Zawilski, S.M., and Cox, E.C. (2005). Real-time kinetics of gene activity in individual bacteria. Cell 123, 1025–1036.

24. Griffiths, R.E., Kupzig, S., Cogan, N., Mankelow, T.J., Betin, V.M.S., Trakarnsanga, K., Massey, E.J., Lane, J.D., Parsons, S.F., and Anstee, D.J. (2012) Maturing reticulocytes internalize plasma membrane in glycophorin A-containing vesicles that fuse with autophagosomes before exocytosis. Blood 119 (26) 6296–6306.

25. Hansen, A.S. and Zechner, C. (2021) Promoters adopt distinct dynamic manifestations depending on transcription factor context. Molecular Systems Biology, 17 (2) e9821.

26. Harper, C.V., Finkenstädt, B., Woodcock, D.J., Friedrichsen, S., Semprini, S., Ashall, L., Spiller, D.G., Mullins, J.J., Rand, D.A., Davis, J.R.E. et al. (2011) Dynamic analysis of stochastic transcription cycles. PLOS Biol. 9 (4), e1000607

27. Hay, D., Hughes, J.R., Babbs, C., Davies, J.O.J., Graham, B.J., Hanssen, L.L.P., Kassouf, M.T., Oudelaar, A.M., Sharpe, J.A., Suciu, M.C., et al. (2016). Genetic dissection of the α-globin super-enhancer in vivo. Nature Genetics 48, 895–903.

28. Heinrich, S., Sidler, C.L., Azzalin, C.M., and Weis, K. (2017) Stem-loop RNA labeling can affect nuclear and cytoplasmic mRNA processing. RNA 23 134–141

29. Jeziorska, D.M., Murray, R.J.S., De Gobbi, M., Gaentzsch, R., Garrick, D., Ayyub, H., Chen, T., Li, E., Telenius, J., Lynch, M., et al. (2017) DNA methylation of intragenic CpG islands depends on their transcriptional activity during differentiation and disease. Proceedings of the National Academy of Sciences 114 (36), E7526–E7535.

30. Keller, G., Kennedy, M., Papayannopoulou, T. and Wiles, M. V. (1993) Hematopoietic commitment during embryonic stem cell differentiation in culture. Molecular and Cellular Biology 13 (1) 473–486.

31. Keller, G.M. (1995). In vitro differentiation of embryonic stem cells. Current Opinion in Cell Biology 7, 862–869.

32. Kingsley, P.D., Greenfest-Allen, E., Frame, J.M., Bushnell, T.P., Malik, J., McGrath, K.E., Stoeckert, C.J., and Palis, J. (2013). Ontogeny of erythroid gene expression. Blood 121, e5–e13.

33. Lee, C., Shin, H., and Kimble, J. (2019). Dynamics of Notch-Dependent Transcriptional Bursting in Its Native Context. Developmental Cell 50, 426–435.e4.

34. Lim, F. and Peabody, D. S. (2002). RNA recognition site of PP7 coat protein. Nucleic Acids Research 30, 4138–4144.

35. Lim, B., Fukaya, T., Heist, T., and Levine, M. (2018). Temporal dynamics of pair-rule stripes in living Drosophila embryos. Proceedings of the National Academy of Sciences 115, 8376–8381.

36. Mei, Y., Liu, Y., and Ji, P. (2021) Understanding terminal erythropoiesis: An update on chromatin condensation, enucleation, and reticulocyte maturation. Blood Reviews 46, 100740

37. Molina, N., Suter, D.M., Cannavo, R., Zoller, B., Gotic, I., and Naef, F. (2013). Stimulus-induced modulation of transcriptional bursting in a single mammalian gene. Proceedings of the National Academy of Sciences 110, 20563–20568.

38. Muramoto, T., Cannon, D., Gierliński, M., Corrigan, A., Barton, G.J., and Chubb, J.R. (2012). Live imaging of nascent RNA dynamics reveals distinct types of transcriptional pulse regulation. Proceedings of the National Academy of Sciences of the United States of America 109, 7350–7355.

39. Nichols, J., Evans, E.P., and Smith, A.G. (1990) Establishment of germ-line-competent embryonic stem (ES) cells using differentiation inhibiting activity. Development 110 (4) 1341–8.

40. Ochiai, H., Sugawara, T., Sakuma, T., Yamamoto, T. (2014) Stochastic promoter activation affects Nanog expression variability in mouse embryonic stem cells. Sci. Adv. 4: 7125.

41. Orkin, S.H., Swan, D., and Leder, P. (1975) Differential expression α-and β-globin genes during differentiation of culture erythroleukaemic cells. J. Biol. Chem. 250 8753–8760.

42. Oudelaar, A.M., Beagrie, R.A., Gosden, M., de Ornellas, S., Georgiades, E., Kerry, J., Hidalgo, D., Carrelha, J., Shivalingam, A., El-Sagheer, A.H., et al. (2020). Dynamics of the 4D genome during in vivo lineage specification and differentiation. Nature Communications 11 2722.

43. Oudelaar, A.M., Beagrie, R.A., Kassouf, M.T., and Higgs, D.R. (2021) The mouse alpha-globin cluster: a paradigm for studying genome regulation and organization. Current Opinion in Genetics and Development

44. Pijuan-Sala, B., Griffiths, J.A., Guibentif, C., Hiscock, T.W., Jawaid, W., Calero-Nieto, F.J., Mulas, C., Ibarra-Soria, X., Tyser, R.C.V., Ho, D.L.L., et al. (2019). A single-cell molecular map of mouse gastrulation and early organogenesis. Nature 566, 490–495.

45. Rodriguez, J., Ren, G., Day, C.R., Zhao, K., Chow, C.C., and Larson, D.R. (2019). Intrinsic Dynamics of a Human Gene Reveal the Basis of Expression Heterogeneity. Cell 176, 213–226.e18.

46. Scholes, C., Biette, K.M., Harden, T.T., and DePace, A.H. (2019). Signal Integration by Shadow Enhancers and Enhancer Duplications Varies across the Drosophila Embryo. Cell Reports 26, 2407–2418.e5.

47. Smith, A.G. (1991) Culture and differentiation of embryonic stem cells. Journal of tissue culture methods 13, 89–94.

48. Socolovsky, M., Nam, H., Fleming, M.D., Haase, V.H., Brugnara, C., and Lodish, H.F. (2001). Ineffective erythropoiesis in Stat5a−/−5b−/− mice due to decreased survival of early erythroblasts. Blood 98, 3261–3273.

49. Stavreva, D.A., Garcia, D.A., Fettweis, G., Gudla, P.R., Zaki, G.F., Soni, V., McGowan, A., Williams, G., Huynh, A., Palangat, M., et al. (2019). Transcriptional Bursting and Co-bursting Regulation by Steroid Hormone Release Pattern and Transcription Factor Mobility. Molecular Cell 75, 1161–1177.e11.

50. Suter, D.M., Molina, N., Gatfield, D., Schneider, K., Schibler, U., and Naef, F. (2011). Mammalian genes are transcribed with widely different bursting kinetics. Science 332, 472–474.

51. Telenius, J., The WIGWAM Consortium, and Hughes, J.R. (2018) NGseqBasic – a single-command UNIX tool for ATAC-seq, DNaseI-seq, Cut-and-Run, and ChIP-seq data mapping, high-resolution visualisation, and quality control. bioRxiv 393413, doi: 10.1101/393413.

52. Tunnacliffe, E., and Chubb, J.R. (2020). What Is a Transcriptional Burst? Trends in Genetics 36, 288–297.

53. Tunnacliffe, E., Corrigan, A.M., and Chubb, J.R. (2018). Promoter-mediated diversification of transcriptional bursting dynamics following gene duplication. Proceedings of the National Academy of Sciences 115, 8364–8369.

54. Tutucci, E., Vera, M., Biswas, J., Garcia, J., Parker, R., and Singer, R.H. (2018) An improved MS2 system for accurate reporting of the mRNA life cycle. Nature Methods 15, 81–89.

55. Waggoner, S. A. and Liebhaber, S.A. (2003) Regulation of alpha-globin mRNA stability. Experimental Biology and Medicine. 228, 387–395.

56. Zhang, J., Socolovsky, M., Gross, A.W., and Lodish, H.F. (2003). Role of Ras signaling in erythroid differentiation of mouse fetal liver cells: functional analysis by a flow cytometry–based novel culture system. Blood 102, 3938–3946.

57. Zoller, B., Nicolas, D., Molina, N., and Naef, F. (2015). Structure of silent transcription intervals and noise characteristics of mammalian genes. Molecular Systems Biology 11, 823–823.

