## Supplementary information for "On-microscope staging of live cells reveals changes in the dynamics of transcriptional bursting during differentiation"

### Supplementary Tables

#### Supplementary Table 1: Primers for RT-qPCR

| Gene | Primer | Sequence |
|---|---|---|
| *Hba-a1/2* | Forward | CTGGGGAAGACAAAAGCAAC |
| | Reverse | GCCGTGGCTTACATCAAAGT |
| *Hba-a1/2* (nascent) | Forward | GTGTGGATCCCGTCAACTTC |
| | Reverse | CCACTATGTTCCCTGCCTTG |
| *Hba-x* | Forward | CTGTCTGCTGGTCACAATGG |
| | Reverse | GGGAGGAGAGGGATCATAGC |
| *Rn18s* | Forward | GTAACCCGTTGAACCCCATT |
| | Reverse | CCATCCAATCGGTAGTAGCG |
| *Tfrc* (CD71) | Forward | TCTGGAATCCCAGCAGTTTC |
| | Reverse | ACCATTTGGTTGAGCTGAGG |
| *Gypa* (Ter119) | Forward | ACTCCTGTGGTGGCTTCAAC |
| | Reverse | TCCTCCAATGTGTGGTGAGA |
| *Gata1* | Forward | CCCAAGAAGCGAATGATTGT |
| | Reverse | TCCGCCAGAGTGTTGTAGTG |
| *Gata2* | Forward | GCAACCCTTACTACGCCAAC |
| | Reverse | GCTGTGCAACAAGTGTGGTC |
| *Flk1* | Forward | GCTTTCGGTAGTGGGATGAA |
| | Reverse | GGCCTTCCATTTCTGTACCA |
| *Tal1* | Forward | TCTGATGGTCCTCACACCAA |
| | Reverse | GTGGGGATCAGCTTTCTGAG |
| *CD44* | Forward | TGGATCCGAATTAGCTGGAC |
| | Reverse | AGCTTTTTCTTCTGCCCACA |
| *Runx1* | Forward | CCAGCCTCTCTGCAGAACTT |
| | Reverse | GACGGCAGAGTAGGGAACTG |
| *Klf2* | Forward | AACTGCGGCAAGACCTACAC |
| | Reverse | TCCTTCCCAGTTGCAATGAT |
| *Nanog* | Forward | CCAAAGGATGAAGTGCAAGC |
| | Reverse | GTGCTGAGCCCTTCTGAATC |

#### Supplementary Note

##### Classification of transcriptional behaviours from time series data

Using summary statistics such as the mean and Fano factor we noticed differences in the distributions of these metrics as cells progress through erythropoiesis. To explore this further we measured the total time spent in the active state (ON duration) for each cell and compared this to the total transcriptional output (defined by the area under the curve between the ON threshold as a lower bound, and the fluctuating spot intensity as an upper bound).

In general, when comparing the relationship between these two variables we observed that the majority of cells are found along a baseline trajectory where total transcriptional output slowly increases as ON duration increases, suggesting these cells are predominantly transcribing at a low, basal rate (Fig. 5A, B). To describe the 'average' relationship between ON duration and transcriptional output we used local regression. This nicely captured this basal behaviour of most cells but at high ON duration the gradient of the regression showed a marked increase, with a clear inflection point at around 53.5 min ON duration (Fig. 5A). This suggested that on average the burst amplitude in near-continuously active cells is higher than those which are actively only intermittently, since the gradient of the regression depends on the relative changes in total transcriptional output (and therefore average burst amplitude) with increased ON duration. Given the fact that Dar et al. (2012) previously described a similar phenomenon where burst size (of which amplitude is a component) is increased only beyond a particular threshold at high levels of expression, we wanted to classify cells according to this change in bursting behavior. Therefore, we used this inflection point to classify cells as either 'high ON' (active > 53.5 min, > 89% of the time) or 'low ON' (active < 53.5 min).

We then wanted to classify cells into those exhibiting largely basal amplitude bursts, and those transcribing with high amplitude bursts, at either low or high ON durations (Fig. 5D, E). To do this, we first extrapolated the trajectory of basal amplitude bursting cells using a simple quadratic fit of the LOWESS regression itself in low ON cells only (Fig. 5C).

We then calculated the residual of each cell to this extrapolated fit to enable quantification of the number of cells which lie significantly above the basal amplitude duration-output trajectory (Fig. 5C). Given that those with higher ON duration have a greater chance of having sufficiently increased transcriptional output to lie above this trajectory (simply due to having been active for longer) we normalized these residuals by the ON duration (Fig. 5C). We then defined a threshold above which cells would be classified as ‘high amplitude’ bursting cells using the distribution of these normalized residuals. Specifically, we defined the threshold as one standard deviation away from the median of normalized residuals at low ON duration (Fig. 5C).

To enable us to check that this analysis was not affected by the empirically defined threshold for when a gene is called as active or ON we repeated this analysis with different values for this threshold (Extended Data Fig. 9B, C). The inflection point was manually identified in each case according to the LOWESS regression, as before. Little change was observed in the relative proportions of cells exhibiting different transcriptional behaviours across the differentiation stages using this approach, suggesting these observations are robust to variation in the spot intensity ON threshold (Extended Data Fig. 9C).

**Benefits of on-microscope staining for staging cells in differentiation**

In this study we exploit a simple on-microscope staining approach (Eilken et al., 2009) to enable us to analyse cells across a range of differentiation states. The benefits of such an approach are several-fold. Firstly, the sensitivity of nascent transcriptional activity to environmental stimuli (Cesbron et al., 2015; Falo-Sanjuan et al., 2019; Kindgren et al., 2019; Mahat et al., 2016; Stavreva et al., 2019) illustrates the need for gentle experimental practices when studying transcription. Popular methods for isolation of immunophenotyped cells such as magnetic bead selection can induce a stress response, with FACS having a somewhat lesser effect (Beliakova-Bethell et al., 2013). On-microscope labelling therefore represents an attractive alternative by reducing the mechanical stimulation of cells. Secondly, given the time required for the above isolation methods (1-4 h depending on the rarity of cell populations) as well as the need to allow cells to acclimatise (30 min-1 h) before imaging nascent transcription, our approach offers significant time savings depending on the number of populations to be imaged. Finally, physiological changes in development, including transcription, frequently occur along a continuum of cell states. In contrast to the other methods mentioned above, our approach enables simultaneous capture of transcription dynamics across this developmental spectrum without the need for arbitrary grouping of cells before the experiment takes place. One limitation of this method is the requirement for stratification of differentiation stages with only a few cell surface markers (given the limits on number of imaging channels for live cell confocal microscopy). However, we believe there are numerous examples of differentiation programmes for which this should not be a hindrance, such as early T cell development (Kernfeld et al., 2018; Shukla et al., 2017), or during reprogramming of human pluripotent stem cells to a naïve state (Bredenkamp et al., 2019).

**References**

Beliakova-Bethell, N., Massanella, M., White, C., Lada, S.M., Du, P., Vaida, F., Blanco, J., Spina, C.A., and Woelk, C.H. (2013). The effect of cell subset isolation method on gene expression in leukocytes. Cytometry Part A *85*, 94–104.

Bredenkamp, N., Stirparo, G.G., Nichols, J., Smith, A., and Guo, G. (2019). The Cell-Surface Marker Sushi Containing Domain 2 Facilitates Establishment of Human Naive Pluripotent Stem Cells. Stem Cell Reports *12*, 1212–1222.

Cesbron, F., Oehler, M., Ha, N., Sancar, G., and Brunner, M. (2015). Transcriptional refractoriness is dependent on core promoter architecture. Nature Communications *6*, 6753.

Dar, R.D., Razooky, B.S., Singh, A., Trimeloni, T.V., McCollum, J.M., Cox, C.D., Simpson, M.L., and Weinberger, L.S. (2012). Transcriptional burst frequency and burst size are equally modulated across the human genome. Proceedings of the National Academy of Sciences *109*, 17454–17459.

Falo-Sanjuan, J., Lammers, N.C., Garcia, H.G., and Bray, S.J. (2019). Enhancer Priming Enables Fast and Sustained Transcriptional Responses to Notch Signaling. Developmental Cell *50*, 411–425.e8.

Kernfeld, E.M., Genga, R.M.J., Neherin, K., Magaletta, M.E., Xu, P., and Maehr, R. (2018). A Single-Cell Transcriptomic Atlas of Thymus Organogenesis Resolves Cell Types and Developmental Maturation. Immunity *48*, 1258-1270.e6.

Kindgren, P., Ivanov, M., and Marquardt, S. (2019). Native elongation transcript sequencing reveals temperature dependent dynamics of nascent RNAPII transcription in Arabidopsis. Nucleic Acids Research *48*, 2332–2347.

Mahat, D.B., Salamanca, H.H., Duarte, F.M., Danko, C.G., and Lis, J.T. (2016). Mammalian Heat Shock Response and Mechanisms Underlying Its Genome-wide Transcriptional Regulation. Molecular Cell *62*, 63–78.

Shukla, S., Langley, M.A., Singh, J., Edgar, J.M., Mohtashami, M., Zúñiga-Pflücker, J.C., and Zandstra, P.W. (2017). Progenitor T-cell differentiation from hematopoietic stem cells using Delta-like-4 and VCAM-1. Nat Methods *14*, 531–538.
